# Exploring the interspecific interactions and the metabolome of the soil isolate *Hylemonella gracilis*

**DOI:** 10.1101/2021.02.11.430889

**Authors:** Olaf Tyc, Purva Kulkarni, Adam Ossowicki, Vittorio Tracanna, Marnix H. Medema, Peter van Baarlen, W.F.J. van IJcken, Koen J. F. Verhoeven, Paolina Garbeva

## Abstract

Microbial community analysis of aquatic environments showed that an important component of its microbial diversity consists of bacteria with cell sizes of ~0.1 μm. Such small bacteria can show genomic reductions and metabolic dependencies with other bacteria. However, so far no study investigated if such bacteria exist in terrestrial environments like e.g. soil.

Here, we isolated soil bacteria that passed through a 0.1 μm filter, by applying a novel isolation and culturing approach. The complete genome of one of the isolates was sequenced and the bacterium was identified as *Hylemonella gracilis*. A set of co-culture assays with phylogenetically distant soil bacteria with different cell and genome sizes was performed. The co-culture assays revealed that *H. gracilis* grows better when interacting with other soil bacteria like *Paenibacillus* sp. AD87 *and Serratia plymuthica*. Transcriptomics and metabolomics showed that *H. gracilis* was able to change gene expression, behavior, and biochemistry of the interacting bacteria without direct cell-cell contact.

Our study indicates that bacteria are present in the soil that can pass through a 0.1 μm filter. These bacteria may have been overlooked in previous research on soil microbial communities. Such small bacteria, exemplified here by *H. gracilis,* are able to induce transcriptional and metabolomic changes in other bacteria upon their interactions in soil. In vitro, the studied interspecific interactions allowed utilization of growth substrates that could not be utilized by monocultures, suggesting that biochemical interactions between substantially different sized soil bacteria may contribute to the symbiosis of soil bacterial communities.

**Importance:** Analysis of aquatic microbial communities revealed that parts of its diversity consist of bacteria with cell sizes of ~0.1 μm. Such bacteria can show genomic reductions and metabolic dependencies with other bacteria. So far, no study investigated if such bacteria exist in terrestrial environments e.g. soil. By applying a novel isolation method, we show that such bacteria also exist in soil. The isolated bacteria was identified as *Hylemonella gracilis*.

Co-culture assays with phylogenetically different soil bacteria revealed that *H. gracilis* grows better when co-cultured with other soil bacteria. Transcriptomics and metabolomics showed that *H. gracilis* was able to change gene expression, behavior, and biochemistry of the interacting bacteria without direct contact. Our study revealed that bacteria are present in soil that can pass through 0.1 μm filters. Such bacteria may have been overlooked in previous research on soil microbial communities and may contribute to the symbiosis of soil bacterial communities.

## Introduction

Bacteria are ubiquitous living organisms with various cell shapes and sizes surrounding us in all environments (1, 2). Soil is the most complex habitat harboring the largest diversity and density of bacteria known to date (cell densities ranging from 10^7^ to 10^10^ cells/g of soil (3–5). Soil bacteria are part of a community where they are in constant interaction with their own and other species (6–8). Bacteria produce and release a plethora of metabolites into their environment. In this way, they not only chemically modify their niche but also affect the behavior and the secondary metabolite production of nearby bacteria (9–11). Soil bacteria are known to produce a wide range of soluble and volatile secondary metabolites with different physicochemical and biological properties (7, 12–14). In contrast to soluble compounds, volatile organic compounds (VOCs) are rather small molecules (< 300 Da) that can diffuse easily through air- and water-filled soil pores (15–17). These physicochemical properties make VOCs ideal metabolites for long-distance communication and interactions between soil microorganisms (18–21).

In aquatic environments, bacteria are naturally found at lower cell densities compared to soil (10^3^ - 10^6^ cells/mL) (22–24). Recent studies have shown that a significant component of aquatic microbial diversity consists of bacteria with small cell sizes of about ~0.1 μm (25–27). However, little is known if bacteria with such cell sizes exist in soil environments, for instance in water-filled soil pores. One can assume that a small cell size can be an advantage in challenging environments like soil. However, the distribution of microorganisms in soil is influenced by its water and moisture content, and a low soil moisture content leads to lower connectivity between soil pores, and thus to a lower number of accessible micro-habitats.

Small bacterial cell size is often linked to a small genome size caused by genome streamlining (28). Recent metagenomics studies suggest that genome streamlining is ubiquitous in bacteria (29, 30). In some cases, the primary metabolism of one organism can be directly built on the primary metabolism of another organism, known as syntrophic relationships (31, 32). The Black Queen Hypothesis states that genome-streamlined organisms have an evolutionary advantage because of the loss of genes whose function can be replaced by bacteria in the surrounding environment, effectively conserving energy (33). Since bacteria with fewer genes have less adaptive capacity compared to bacteria with more genes, many of them are expected to depend on specific environmental conditions or on the presence of other specific organisms (34) to produce metabolites that support their persistence.

Here, we aimed to explore if bacteria that are able to pass through 0.1 μm filters are present in soil, and if such bacteria are cultivable. We further investigated their interaction with phylogenetically different bacteria commonly occurring in soil. The major research questions were if, and how inter-specific interactions between bacteria that pass a 0.1 μm filter and other common soil bacteria that cannot pass 0.1 μM filters affected their fitness, behavior, gene expression, and the production of secondary metabolites.

## Materials and Methods

### Isolation and identification of bacteria that pass through 0.1 μm filters Isolation of *H. gracilis* from soil

After removing the grassland vegetation, a topsoil core was collected and mixed, a sample of 10 g was suspended in 90 ml of 10 mM Phosphate-buffer (pH 6.5) and shaken with gravel (2-4 mm) at 250 rpm for 45 min. The extract was filtered through sterile gauze pads and subsequently through sterile Whatmann 1mm paper filter using Buchner funnel. Purified extract was filtered again through a syringe filter 0.2 μM and afterwards through 0.1 μM filter (GE-Healthcare). The filtered extract was plated on 1/10th TSBA plates immediately after isolation and incubated at 25 °C (**Supplementary Figure 1**). The plates were inspected daily using a stereomicroscope (Leica M205C) screening for bacteria colonies.

### Bacteria and culture conditions

The bacterial strains used in this study are the Gram-negative strain *S. plymuthica* PRI-2C (gamma-Proteobacteria) (35), the Gram-positive strain *Paenibacillus* sp. AD87 (Firmicutes) (10, 36, 37) and the Gram-negative *H. gracilis* isolate NS1 (beta-Proteobacteria). The bacterial isolates were pre-cultured from −80 °C glycerol stocks on 1/10^th^ TSBA (38) or on LB-A plates (*H. gracilis*) (LB-Medium Lennox, Carl Roth GmbH + Co. KG, 20 gL^-1^ Bacto Agar) and incubated at 24 °C prior application. All bacterial isolates are listed in **Supplementary Table 1.**

### Identification of *H. gracilis*

For the identification of *H. gracilis* 16S rRNA PCR was performed from grown colonies in a 50 μl PCR-GoTaq™ green master mix (Promega Corp. Madison, USA cat# M712). For 16S rRNA gene amplification the following primers were used: forward primer 27f (5’-AGA GTTT GAT CMT GGC TCAG −3’), reverse primer 1492r (5’-GRT ACC TTG TTA CGA CTT −3’), amplifying ~1465 bp from the 16S rRNA gene (39, 40) (modified). All PCR reactions were performed on a BIO-RAD C1000 Touch™ PCR machine (BIO-RAD, Veenendaal, the Netherlands) with these settings: initial cycle 95 °C for 3 min. and 30 cycles of 94 °C for 30 sec., 55 °C for 45 sec. and 72 °C for 1 min. and final extension at 72 °C for 5 minutes. The PCR products were purified using the Qiagen PCR purification kit and sent to MACROGEN (MACROGEN Europe, Amsterdam, the Netherlands for 16S rRNA sequencing.

### Microscopy

Microscopy pictures of *H. gracilis* cells were taken at 400-fold magnification with an Axio Imager M1 microscope (Carl Zeiss, Germany) under phase-contrast illumination with an AxioCam MRm camera. Macroscopic colony pictures of *H. gracilis* were taken with an OLYMPUS Binocular at 20 X magnification. Images were analyzed with AXIO VISION v4.7 (Carl Zeiss Imaging Solutions GmbH, Germany).

### Bacterial interactions assays

After four-days of pre-culture, a single colony of *Paenibacillus* sp. AD87 and *S. plymuthica* PRI-2C and *H. gracilis* was picked and inoculated in 20 mL 1/10^th^ TSB *(Paenibacillus* sp. AD87 and *S. plymuthica* PRI-2C) and grown overnight at 24 °C at 220 rpm. For the inoculation of *H. gracilis* a single colony was picked from a TSBA plate and inoculated in 20 mL LB-medium and grown for 3 days at 24 °C, 200 rpm. For the interaction assay an inoculation mix of each bacterial strain *(Paenibacillus* sp. AD87, *S. plymuthica* PRI-2C, *H. gracilis*) was prepared by diluting the bacterial isolates in 20 mL of 10 mM Phosphate-buffer (pH 6.5) to an OD_600_ of 0.005 *(Paenibacillus* sp. AD87 and *S. plymuthica* PRI-2C) or to an OD_600_ of 0.05 (*H. gracilis*), which corresponds to 10^5 CFU/mL. A droplet of 10 μl was added in the middle of a 6 cm diameter Petri dish (monocultures) or next to each other in a distance of ~0.5 cm (pairwise interactions). All treatments were performed in triplicates on 1/10^th^ TSBA plates incubated at 24 °C. After the growth time of 3 days the bacteria were scratched and washed from the plates by using sterile cell scratchers and Phosphate buffer. For the enumeration of the cell counts (CFU/mL) dilution series of the scratched bacteria were prepared and plated in triplicates on 1/10^th^ TSBA plates and grown for 48 hours. Enumeration was carried out on an aCOlyte Colony Counter (Don Whitley Scientific, Meintrup DWS Laborgeräte GmbH, Germany).

### Enumeration of growth inhibitory or growth promoting effects of cell-free supernatants of *Paenibacillus* sp. AD87and *S. plymuthica* PRI-2C on the growth of *H. gracilis*

A bacterial growth assay in liquid media supplemented with cell-free supernatants (CFS) of *Paenibacillus* sp. AD87 and *S. plymuthica* PRI-2C was conducted. For the assay single colonies of *Paenibacillus* sp. AD87, *S. plymuthica* PRI-2C and *H. gracilis* were inoculated in 20 mL 1/10^th^ TSB (*Paenibacillus* sp. AD87 and *S. plymuthica* PRI-2C) or in 20 mL LB-medium (*H. gracilis*) and grown overnight at 24 °C, 220 rpm or for three days (*H. gracilis*). For the preparation of *Paenibacillus* sp. AD87 and *S. plymuthica* PRI-2C cell-free supernatant (CFS) the grown cultures were centrifuged at 5000 rpm for 20 minutes (at room temperature) and filtered through 0.2 μM filters (GE Healthcare). For the assay, *H. gracilis* was inoculated into 20 mL liquid LB-media at an OD_600_ of 0.05. The growth media was then supplemented either with 20 % (v/v) CFS of *Paenibacillus* sp. AD87 or *S. plymuthica* PRI-2C or with 20 % (v/v) of filter sterilized liquid 1/10^th^ TSB media (control). The cultures were incubated at 24 °C at 220 rpm for 7 days and the bacterial growth was monitored by optical density (absorbance at 600nm) measurements and by plate counting. After five days of growth, the CFU/ml of *H. gracilis* grown in presence of CFS of *Paenibacillus* sp. AD87 or *S. plymuthica* PRI-2C were enumerated by plate counting. For this, the cultures were sampled and dilution series were prepared in triplicates and a volume of 100 μl of each serial dilution was plated in three replicates with a disposable Drigalski spatula on 1/10^th^ TSBA plates. CFU Enumeration was carried out on an aCOlyte Colony Counter (Don Whitley Scientific, Meintrup DWS Laborgeräte GmbH, Germany).

### DNA isolation and genome sequencing of *H. gracilis*

Genomic DNA of *H. gracilis* was extracted using a QIAGEN Genomic-tip 500/G DNA kit Qiagen, cat# 10262. Genome sequencing was performed on the PacBio RS II platform (Pacific Biosciences, Menlo Park, CA, USA) using P6-C4 chemistry at the Institute for Genome Sciences (IGS), Baltimore, Maryland, USA. The sequencing resulted in a total of 70,101 reads with N50 of 17 309 nucleotides. The PacBio raw sequences were analyzed using SMRT portal V2.3.0.140936 p.4150482. Sequences were assembled *de novo* with the RS_HGAP_assembly 3 software (Pacific Biosciences, Menlo Park, CA, USA) with default settings on an estimated genome size of 3.8 Mbp. The resulting assemblies were subjected to scaffolding using the RS_AHA_scaffolding 1 software. The genome assembly properties are shown in **Table 1**. Final contigs were annotated using PROKKA V1.11 (41) and InterproScan 5.16 55.0 (42). The whole genome sequence was submitted as *Hylemonella gracilis* strain NS1 to NCBI GenBank (https://www.ncbi.nlm.nih.gov/genbank/) under accession # CP031395.

**Table 1:** Genome assembly statistics and outcome of *in silico* analysis of secondary metabolite gene clusters of *H. gracilis, S. plymuthica* PRI-2C and *Paenibacillus* sp. AD87.

| Feature / Organism | *Hylemonella gracilis* | *S. plymuthica* PRI-2C | *Paenibacillus* sp. AD87 |
| --- | --- | --- | --- |
| contigs | 1 | 1 | 30 |
| bases | 3822245 | 5474685 | 7086713 |
| number of chromosomes | 1 | 1 | 1 |
| size chromosome 1 | 3822245 | 5464425 | 7086713 |
| CDS | 3648 | 4929 | 6216 |
| GC- content (%) | 65.1 | 55.7 | 46.2 |
| number of RNAs | 53 | 109 | 146 |
| genes | 3625 | 5284 | 6375 |
| in silico detected secondary metabolite clusters (antiSMASH) | 3 | 9 | 10 |
| **Total genome size (bases)** | **3822245** | **5474685** | **7086713** |

### *In silico* analysis of secondary metabolite gene clusters

For *in silico* analysis of secondary metabolite gene clusters, the genome sequences of *H. gracilis, Paenibacillus* sp. AD87 and *S. plymuthica* PRI-2C were submitted to the antiSMASH web server (http://antismash.secondarymetabolites.org/) version 4.0 (43).

### RNA isolation and sequencing

Sampling for total RNA extractions was performed in triplicates after five and ten days of incubation on bacteria grown on 1/10^th^ TSBA plates either in co-culture or monoculture as described previously (Bacterial interactions assays on 1/10th TSBA plates). For the isolation of bacterial cell material a volume of 1 mL of 10 mM phosphate buffer (pH 6.5) was added to the surface of the 1/10^th^ TSBA plates and grown bacterial cells were suspended from the plate surface with a disposable cell scratcher (VWR international B.V., the Netherlands). For total RNA extraction the obtained cell suspension was transferred to a tube containing RNA Protect Bacteria Reagent (Qiagen, cat# 76506) and centrifuged for 20 min. at 20,000g, 4 °C. The supernatant was discarded and the resulting cell pellets were stored at −80 °C. Total RNA was extracted using the Aurum Total RNA Mini Kit (BIO-RAD) according to the manufacturer’s protocol. Samples were treated with TURBO DNA free Kit (AMBION) according to the manufacturer’s protocol. The RNA concentration and quality was checked on a NanoDrop Spectrophotometer (ND 2000, Thermo Fisher Scientific, the Netherlands) and on a 1.0 % TBE agarose gel. Samples were subjected to RNA sequencing at the Erasmus Center for Biomics (www.biomics.nl), Erasmus MC, Rotterdam, The Netherlands using the Illumina HiSeq 2500 sequencing platform. The obtained reads were checked for quality using Fastq. For the estimation of the transcripts, the filtered sequences were aligned against the cDNA sequences of *H. gracilis*, *Paenibacillus* sp. AD87 and *S. plymuthica* PRI-2C using Bowtie 2 (2.2.5) (44) with the following settings: -- no-mixed -- no-discordant -- gbar 1000 – end-to-end. Transcript abundance was calculated using RSEM V1.1.26 (45) and differential expression between the treatments was calculated using edgeR V3.2 package in the R environment (46–48).

### Pathway annotations

Please see Supplementary Methods.

### Exploration of missing genes and genome streamlining in *Hylemonella*

RAST annotations of *S. plymuthica* PRI-2C, *Paenibacillus* sp. AD87 and *H. gracilis* were used to compare their genomes and to explore the genomes for missing genes in metabolic pathways (http://rast.nmpdr.org) (49–51). The missing gene sequences were extracted and assigned with KEGG Orthology (52, 53). Presence/absence of genes belonging to metabolic pathways was compared across the three genomes to identify shared genes and pathways and to determine incomplete metabolic pathways in *H. gracilis*.

### Catabolic profiling

To determine the carbon source usage abilities of *H. gracilis* and *Paenibacillus* sp. AD87and *S. plymuthica* PRI-2C strains, Biolog EcoPlate (Labconsult S.A.-N.V., Bruxelles, Belgium) assays were performed (54, 55). Bacteria were cultured in monoculture or in co-culture in single wells of the Biolog EcoPlate™. A, single colony of each bacterial strain was picked and inoculated in 15 mL 1/10^th^ TSB or 15 mL LB-liquid media. Bacteria were grown overnight (*Paenibacillus* sp. AD87 and *S. plymuthica* PRI-2C) or for two days (*H. gracilis*) at 24 °C at 250 rpm. Grown bacteria cultures were washed twice by centrifugation at 4.500 rpm for 15 minutes at room temperature, the supernatant discarded and the pellet was washed and re-suspended in 5 mL of 10 mM phosphate buffer. The re-suspended cultures were diluted to an OD_600_ of 0.005 in 20 mL of 10 mM phosphate buffer either in monoculture or in coculture. For the experiment Biolog EcoPlate™ were inoculated with 100 μl of each bacterial inoculation suspension (monocultures or co-cultures) in each well. For each bacterial monoculture one Biolog EcoPlate™ was inoculated, as well for each co-cultivation pair (*S. plymuthica* PRI-2C with *H. gracilis* and *Paenibacillus* sp. AD87with *H. gracilis)*. Plates were incubated for 1 week and absorbance was measured at 590 nm every 24 hours on a BIOTEK plate reader to determine the ability of the bacterial cultures to use the carbon sources present in the wells.

### Trapping of volatile organic compounds and GC-Q-TOF analysis

Please see Supplementary Methods.

### Ambient mass-spectrometry imaging LAESI-MS data analysis

For LAESI-MS analysis a single colony of each bacterial isolate was picked and inoculated in 20 mL 1/10^th^ TSB *(Paenibacillus* sp. AD87 and *S. plymuthica* PRI-2C) and grown overnight at 24 °C at 220 rpm. For the inoculation of *H. gracilis* a single colony was picked from plate and inoculated in 20 mL LB-medium and grown for three days at 24 °C at 200 rpm. The inoculation mix was prepared by diluting the bacterial isolates in 20 mL of 10 mM Phosphate-buffer (pH 6.5) to an OD_600_ of 0.005. The inoculum mix was pulse-vortexed for 30 sec. and a droplet of 10 μl was added in the middle of a 6 cm diameter Petri dish (monocultures) or next to each other in a distance of approx. 0.5 cm (pairwise interactions). All treatments were inoculated in triplicates on 1/10^th^ TSBA and incubated at 24 °C for five and ten days. After five and ten days of incubation bacterial colonies were cut out of the agar (size approximately 1 – 3 cm^2^) and subjected to LAESI-MS measurement. The LAESI-MS analysis was carried out on a Protea Biosciences DP-1000 LAESI system (Protea Bioscience Inc., Morgantown, WV, USA) coupled to a Waters model Synapt G2S (Waters Corporation, Milford, MA, USA) mass spectrometer. The LAESI system was equipped with a 2940-nm mid-infrared laser yielding a spot size of 100 μm. The laser was set to fire 10 times per x-y location (spot) at a frequency of 10 Hz and 100% output energy. A syringe pump was delivering the solvent mixture of methanol/water/formic-acid (50:50:0.1% v/v) at 2 μL/min to a PicoTip (5cm x 100 μm diameter) stainless steel nanospray emitter operating in positive ion mode at 4000 V. The LAESI was operated using LAESI Desktop Software V2.0.1.3 (Protea Biosciences Inc., Morgantown, WV, USA). The Time of Flight (TOF) mass analyzer of the Synapt G2S was operated in the V-reflectron mode at a mass resolution of 18.000 to 20.000. The source temperature was 150 °C, and the sampling cone voltage was 30 V. The positive ions were acquired in a mass range of 50 to 1200 *m/z*. The MS data was lock mass corrected post data acquisition using leucine encephalin (C_25_H_3_7N_5_O7*m/z*= 556.2771), which was used as an internal standard. All the acquired Waters *.RAW data files were converted to open file format *.imzML using an in-house script written in R. Later, this data was pre-processed in multiple steps to remove noise and to make the data comparable. First, square root transformation was applied to the data to stabilize the variance. Then, baseline correction was performed to enhance the contrast of peaks to the baseline. For better comparison of intensity values and to remove small batch effects, Total-Ion-Current (TIC)-based normalization was applied. This was followed by spectral alignment and peak detection to extract a list of significant mass features for each sample replicate per treatment. In the end, a mass feature matrix was generated with sample replicates for each treatment in columns and mass features in rows. This feature matrix was used to perform further statistical analysis. The preprocessing and peak-detection steps were applied using R scripts developed in-house and using functions available within the MALDIquant R package (56). To perform multivariate analysis, the feature matrix was imported into the online version of Metaboanalyst 4.0 (57). Ion intensity maps displaying the spatial distribution for statistically significant mass features were created using R. Before generating the ion maps, the intensity values for the selected mass features were normalized to the maximum intensity within the image, measured for each mass value individually. Venn diagrams displaying unique and common masses amongst different treatments were drawn using the jvenn tool (58).

## Results

### Isolation and identification of bacteria that pass through 0.1 μM filter

Using a novel bacterial isolation and culture approach, we isolated bacteria from a terrestrial soil sample that were able to pass through 0.22 μm and 0.1 μm pore-size filters. After several days of incubation, only one type of bacterial colonies was observed on the inoculated plates. The grown colonies were identified as *Hylemonella gracilis* (Gram-negative, class betaproteobacteria, order Burkholderiales) by 16S rRNA sequence analysis.

The colonies showed a round and colorless morphology when grown on 1/10^th^ TSBA plates (**Fig. 1a**). Microscopically the bacteria had a spiraled morphology with a length of approximately 6 – 12 μm, which is typical for *Hylemonella* species (**Fig. 1b**).

**Figure 1:**
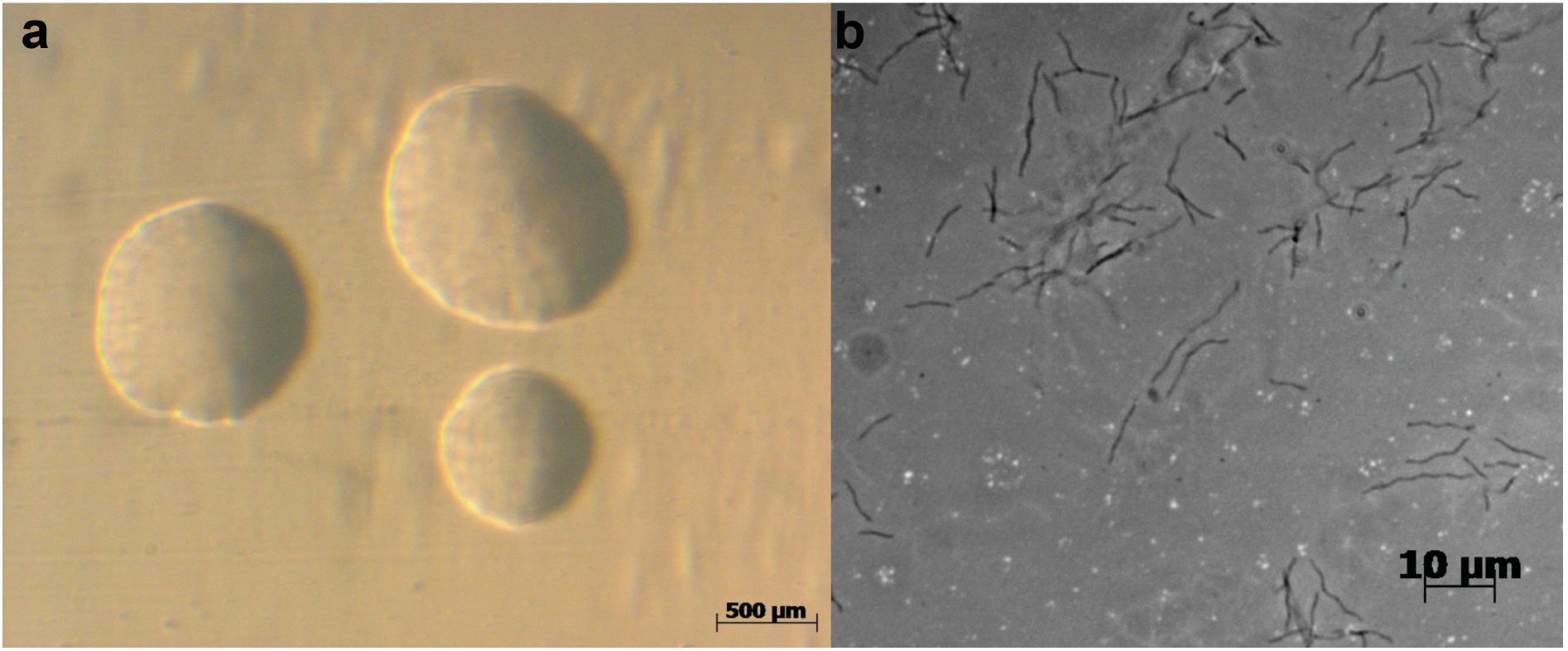
Morphology of *Hylemonella gracilis*. (**a**) on 1/10^th^ TSB-agar plates captured at 20x magnification and (**b**) single bacterial cells captured at 400 X magnification showing their very thin, long and slender appearance in liquid media.

### *Hylemonella* grows better in interaction with other bacteria

To test the hypothesis that small bacteria grow better in presence of normal-sized bacteria, growth of *H. gracilis* was determined in co-culture with two phylogenetically distantly related soil bacteria (*Paenibacillus* sp. AD87 and *S. plymuthica* PRI-2C) and compared to that of the monoculture. The bacterial colony forming units of *H. gracilis* (CFU/mL) obtained on 1/10th TSBA plates from monocultures and co-cultures are summarized in **Fig. 2**. Cell counts of *Paenibacillus* sp. AD87 were 7.68 x 10^7^ CFU/mL in co-culture with *H. gracilis* (**Fig. 2a**). During the interaction with *H. gracilis*, the growth of *S. plymuthica* was significantly negatively affected *(P=0.037)* after five days of incubation by reaching 1.47 x 10^9^ CFU/mL compared to the monocultures (**Fig. 2a**).

**Figure 2:**
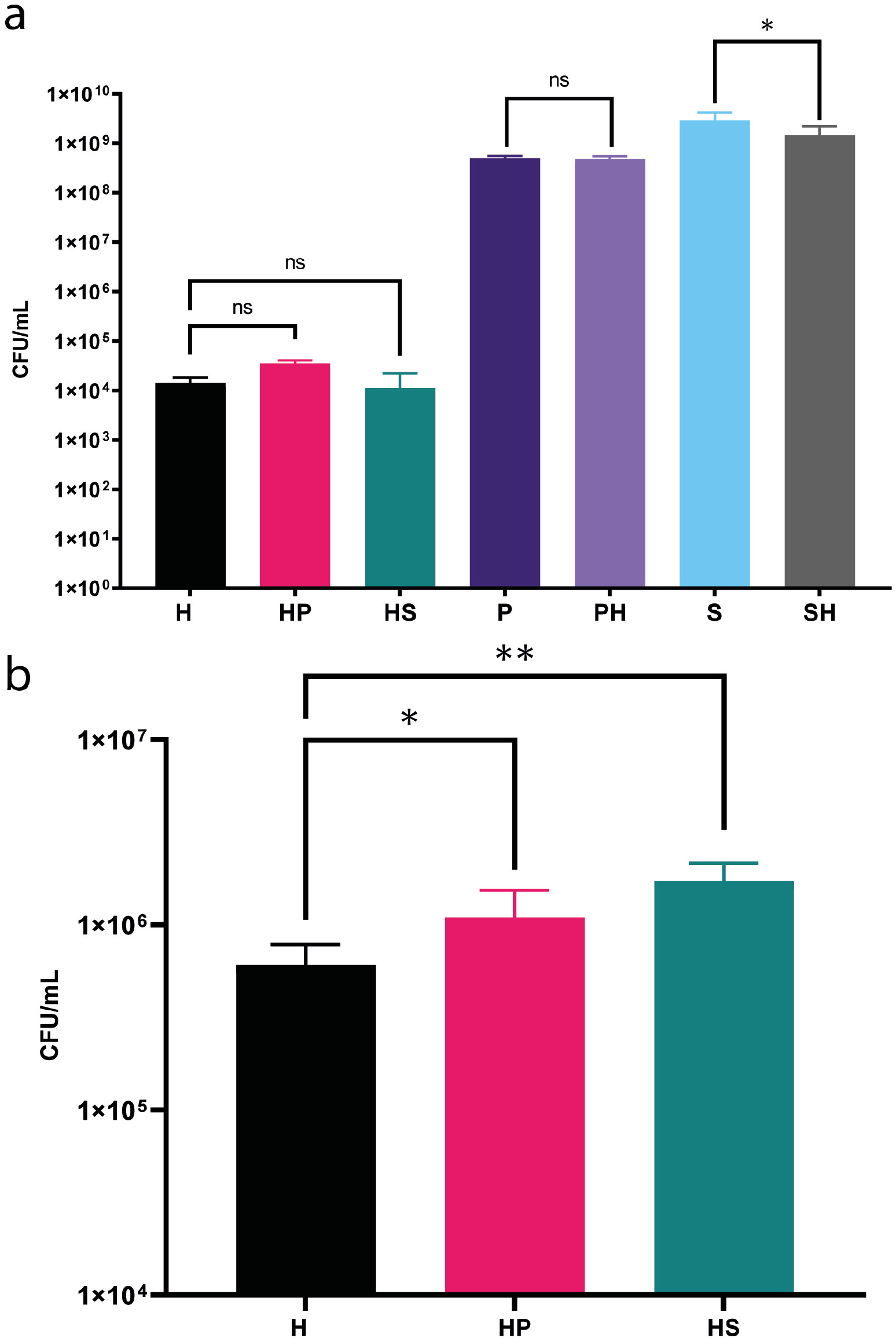
Growth of bacterial mono and cocultures (a) on the plate-based experiment and (b) during the cell-free-supernatant (CFS) experiment. Abbreviations: *H. gracilis* monoculture (H), *Paenibacillus* sp. AD87 monoculture (P), *Paenibacillus* sp. AD87 – *H. gracilis* coculture (HP), *S. plymuthica* PRI-2C monoculture (S)*, S. plymuthica* – *H. gracilis* coculture (SH). *H. gracilis* – *Paenibacillus* sp. AD87 coculture (HP), *H. gracilis* – *S. plymuthica* PRI-2C coculture (HS). Significant differences in colony forming units per milliliter (CFU/mL) between co-cultures (treatment) and monocultures (controls) are indicated by asterisks (ONE-WAY ANOVA, post-hoc TUKEY test).

The bacterial colony forming units (CFU) obtained from *H. gracilis* grown in presence of cell free supernatants (CFS) of *Paenibacillus* sp. AD87 and of *S. plymuthica* are summarized in **Fig. 2b**. *H. gracilis* growth was significantly increased (*P*=0.011) when growing in presence of cell free supernatants of *Paenibacillus* sp. AD87 resulting in higher *H. gracilis* cell counts compared to the monoculture by reaching 1.10 x 10^6^ CFU/mL. In the presence of cell free supernatant from *S. plymuthica* PRI-2C*, H. gracilis* reached the highest cell counts at 1.72 x 10^6^ CFU/mL (*P*=0.000) after five days of incubation (**Fig. 2b**).

### Interspecific interaction between bacterial species allows use of additional substrates

During physiological or catabolic profiling, the metabolism of 31 carbon sources during bacterial growth are measured in 96-wells plates. The catabolic profiling assays revealed that *Paenibacillus* sp. AD87was able to utilize 11 out of the 31 carbon sources in monoculture, while *S. plymuthica* PRI-2C and *H. gracilis* were able to utilize 17 and 16 carbon sources, respectively. Interestingly, three compounds could be utilized only during the co-cultivation of *H. gracilis* with one of the other species, these compounds could not be utilized by any of the species in monoculture. Specifically, alpha-cyclodextrin was utilized only during the cocultivation of *H. gracilis* with *Paenibacillus* sp. AD87, while L-threonine and glycyl-L-glutamic acid were utilized only during the co-cultivation of *S. plymuthica* PRI-2C and *H. gracilis* (**Fig. 3**).

**Figure 3:**
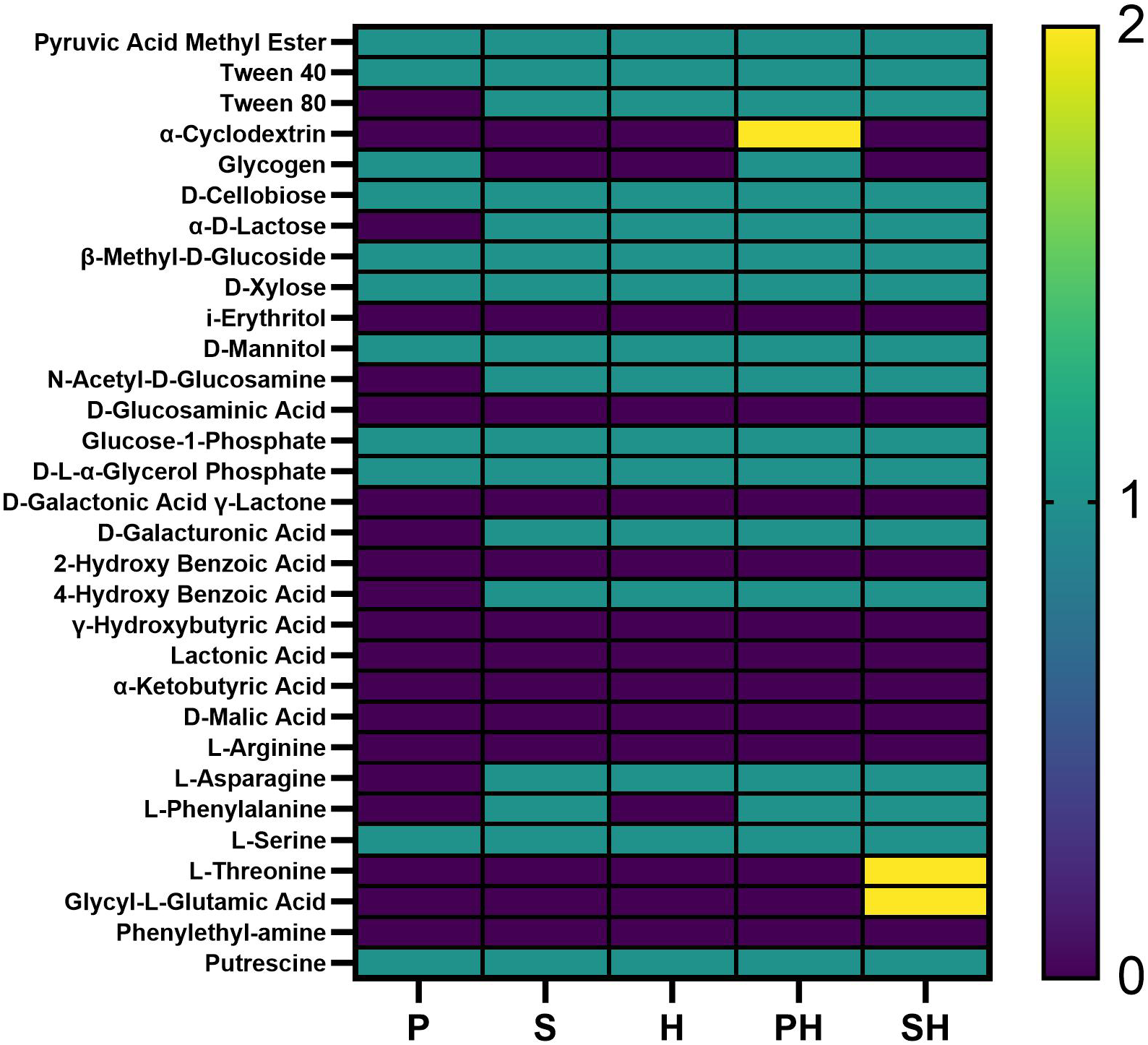
Results of the Biolog EcoPlate™ experiment. Bacteria were inoculated in monoculture or in pairwise combinations on the EcoPlate™ with 31 different carbon sources. Colour code: turquoise=carbon source could be utilized in monoculture (1), yellow= carbon source could be utilized only in co-culture (2) purple= carbon source could not be utilized (0). Abbreviations: *Paenibacillus* sp. AD87 monoculture (P), *Paenibacillus* sp. AD87 – *H. gracilis* coculture, (PH) *H. gracilis* monoculture (H), *S. plymuthica* PRI-2C monoculture (S), *S. plymuthica* PRI-2C – *H. gracilis* coculture (SH).

### Genomic features of *H. gracilis, S. plymuthica* PRI-2C *and Paenibacillus* sp. AD87

Sequencing of the complete genome of *H. gracilis* resulted in a genome size of 3.82 Mbp with 3,648 coding sequences (CDS). As expected, the genome analysis revealed that the genome of *H. gracilis* is smaller and contains fewer genes compared to *S. plymuthica* PRI-2C (5.4 Mbp) and *Paenibacillus* sp. AD87 (7.0 Mbp). The genome features of all three bacteria are summarized in **Table 1**.

### *In silico* analysis of gene clusters encoding for secondary metabolites

The *in silico* tool *antiSMASH* allows the rapid genome-wide identification, annotation and analysis of secondary metabolite biosynthesis gene clusters in bacterial and fungal genomes (59). *In silico* analysis of *Paenibacillus* sp. AD87 revealed a total of 10 gene clusters coding for secondary metabolites. From which two gene clusters encode pathways for producing terpenes, one for bacteriocins, one for lasso peptides, two for lanthipeptides, one for nonribosomal peptides (NRPs), one for others, one for polyketides (type III enzyme mechanism) and one gene cluster for non-NRP Siderophores (**Fig. 4a**). For *S. plymuthica* PRI-2C, nine gene clusters, were found of which two gene clusters were annotated to encode the production of NRPs, one of homoserine lactones, one of aryl polyenes and/or non-NRP Siderophores, one of hybrid polyketide-NRP metabolites, one of thiopeptides, one of butyrolactones, one of terpenes and one of others (**Fig. 4b**). For *H. gracilis* the AntiSMASH analysis revealed that *H. gracilis* possesses relatively few gene clusters related to secondary metabolism. A total of three gene clusters for *H. gracilis* were detected, of which one belonged to the class of bacteriocins, one to the class of terpenes, and one to aryl polyenes, the latter being a homolog to the aryl polyene gene cluster from *Xenorhabdus doucetiae* (Genbank: NZ_FO704550.1) (**Fig. 4c**)

**Figure 4:**
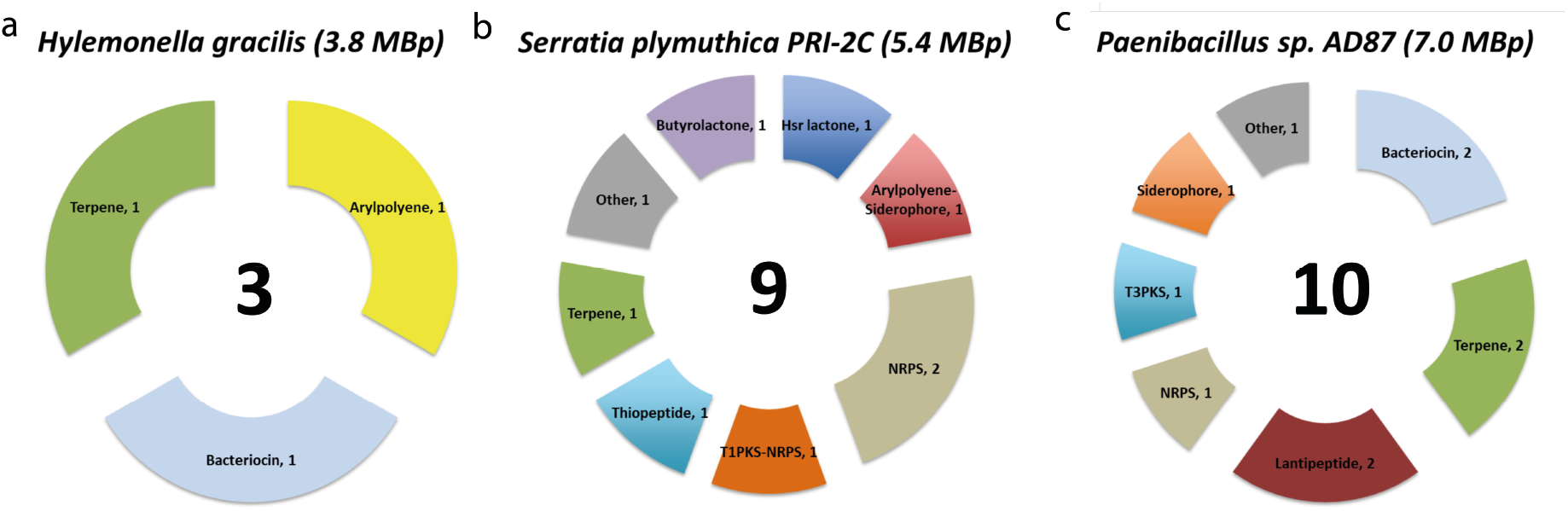
*In silico* comparison of biosynthetic gene clusters (BGCs) predicted by antiSMASH in the genomes of three soil bacteria. From left to right (**a**) *H. gracilis* with a genome size of 3.8 MBp, n= 3 gene clusters for secondary metabolites, (**b**) *S. plymuthica* PRI-2C with a genome size of 5.4 MBp, n=9 gene clusters for secondary metabolites and (**c**) *Paenibacillus* sp. AD87 with a genome size of 7.0 MBp, n=10 gene clusters for secondary metabolites.

### Pathway analysis in *H. gracilis* compared to *S. plymuthica* PRI-2C and *Paenibacillus* sp. AD87

For annotation and Pathway analysis RAST (Rapid Annotation using Subsystem Technology) and OrthoFinder were used. The RAST comparison of *Paenibacillus* sp. AD87 and *H. gracilis* revealed 504 unique enzymes (according to their EC numbers) exclusive for *Paenibacillus* sp. AD87, while 434 were present only in *H. gracilis* and 532 EC numbers were shared by both genomes (**Fig. 5a**). The RAST comparison of *S. plymuthica* PRI-2C and *H. gracilis* revealed that 751 enzymes were present only in *S. plymuthica* PRI-2C, and 260 were present only in *H. gracilis*. 727 EC numbers participating in diverse metabolic pathways were found in both genomes (**Fig. 5b**).

**Figure 5:**
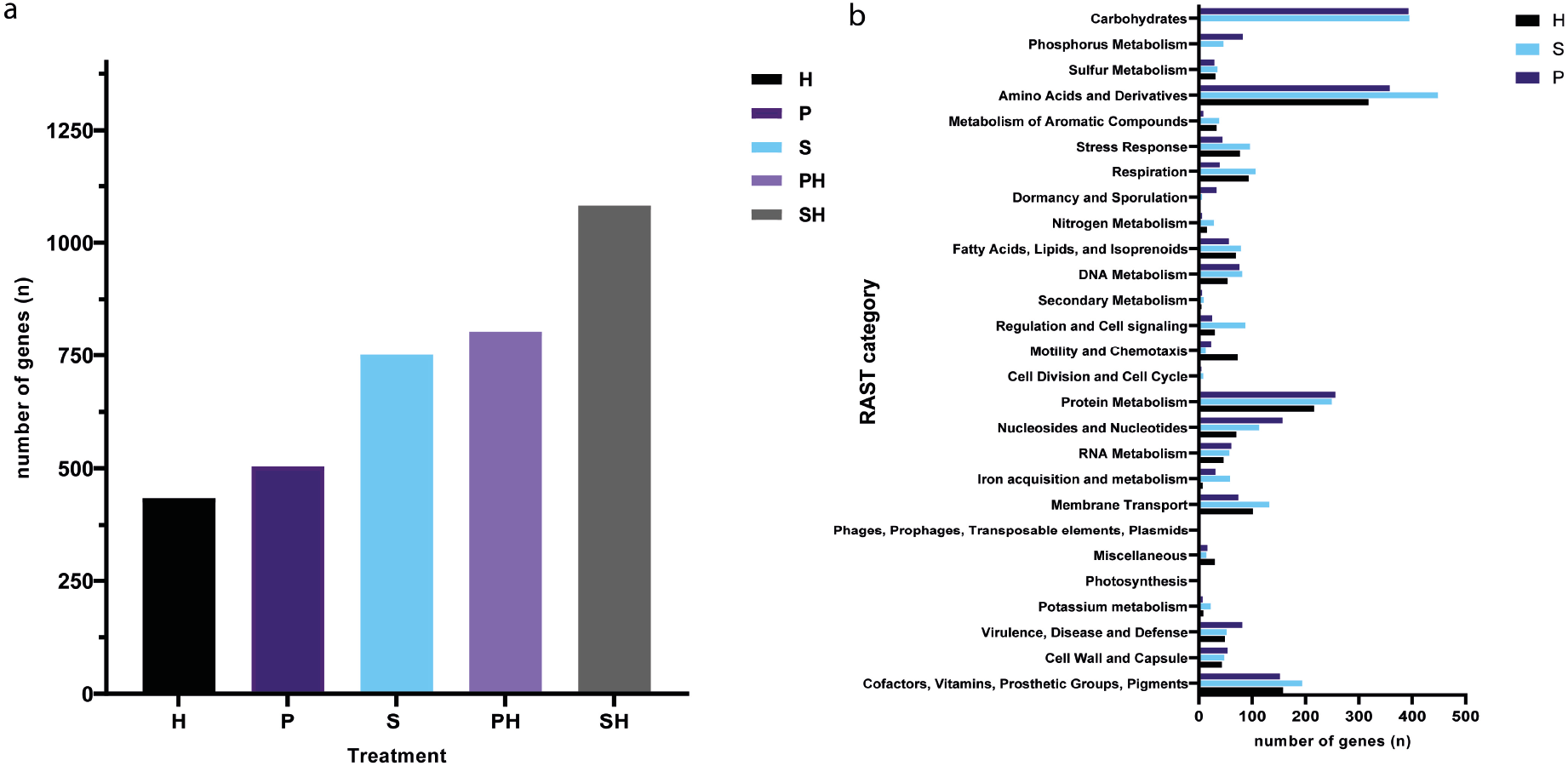
Gene content comparison. (**a**) Box-plot showing the number (n) of all expressed genes (n) for the monocultures of *H. gracilis* (H), *Paenibacillus* sp. AD87 monoculture (P) and *S. plymuthica* PRI-2C monoculture (S) and during the interaction of *H. gracilis* with *Paenibacillus* sp. AD87 (PH) and for the interaction of *H. gracilis* with *S. plymuthica* PRI-2C (SH) determined by RAST. (**b**) Boxplot showing number (n) of expressed genes present in each RAST subsystem category for each of the monocultures.

The missing genes and pathways found by OrthoFinder and EggNOG were annotated with GO terms. The analysis revealed that five genes related to metabolic pathways were absent in *H. gracilis*. Those missing genes were annotated with the following molecular function ontology terms: GO:0008473 (ornithine cyclodeaminase activity), GO:0008696 (4-amino-4-deoxychorismatelyase activity), GO:0003920 (GMP reductase activity), GO:0004035 (alkaline phosphatase activity) and GO:0008442 (3-hydroxyisobutyrate dehydrogenase). We verified if the absence of these molecular functions would render specific pathways obsolete or unavailable in *H. gracilis*. However, alternative pathways routes are present for these genes encoding certain molecular functions according to KEGG database annotations. The pathway analysis by RAST did not reveal the absence of essential genes in *H. gracilis*. Still, the comparison of the number (n) of genes present in each bacteria revealed major differences in several pathways, specifically in the categories “Carbohydrates metabolism” and “Phosphorus metabolism” (**Fig. 5b**). Interestingly, *H. gracilis* possesses no genes for those categories according to RAST, whereas *Paenibacillus* sp. AD87possesses 393 and 82 genes, and *S. plymuthica* PRI-2C 395 and 46 genes, respectively. A major difference in the absolute number of genes in a category is also observed for Amino Acids and Derivatives, for which *H. gracilis* possesses 318 genes, *Paenibacillus* sp. AD87 possesses 358 and *S. plymuthica* PRI-2C 448 genes.

### Effect of interspecific interactions on gene expression

The Transcriptome analysis of monocultures and co-cultures revealed a total of 277 significant differentially expressed genes; where from a total of 100 genes were down-regulated and 177 genes were up-regulated between the different treatments (**Table 2**).

**Table 2:** Overview of the transcriptome analysis, number (n) of significantly differentially expressed genes of *H. gracilis* responding to *S. plymuthica* PRI-2C or to *Paenibacillus* sp. AD87, *Serratia plymuthica* PRI-2C responding to *H. gracilis* and *Paenibacillus* sp. AD87 responding to *H. gracilis* at day 5 and day 10.

| Organism | Interacting organism | time point (d) | Significantly differentially expressed genes |  | Total (n) |
| --- | --- | --- | --- | --- | --- |
|  |  |  | up-regulated | down-regulated |  |
| *Serratia plymuthica* PRI-2C | *Hylemonella gracilis* | 5 | 25 | 36 | 61 |
|  |  | 10 | 10 | 0 | 10 |
| *Paenibacillus* sp. AD87 |  | 5 | 8 | 0 | 8 |
|  |  | 10 | 0 | 0 | 0 |
| *Hylemonella gracilis* | *Serratia plymuthica* PRI-2C | 5 | 1 | 0 | 1 |
|  |  | 10 | 129 | 53 | 182 |
|  | *Paenibacillus* sp. AD87 | 5 | 0 | 0 | 0 |
|  |  | 10 | 4 | 11 | 15 |
| Total (n) |  |  | 177 | 100 | 277 |

#### *Effect of inter-specific interactions on gene expression in Paenibacillus* sp. AD87 *and H. gracilis*

Genes related signal transduction (T) were the category with the most differentially expressed genes during the co-cultivation of *H. gracilis* with *Paenibacillus* sp. AD87 compared to the monoculture of *H. gracilis* (**Supplementary Table 7** and **8**, **Fig. 6a, b**).

**Figure 6:**
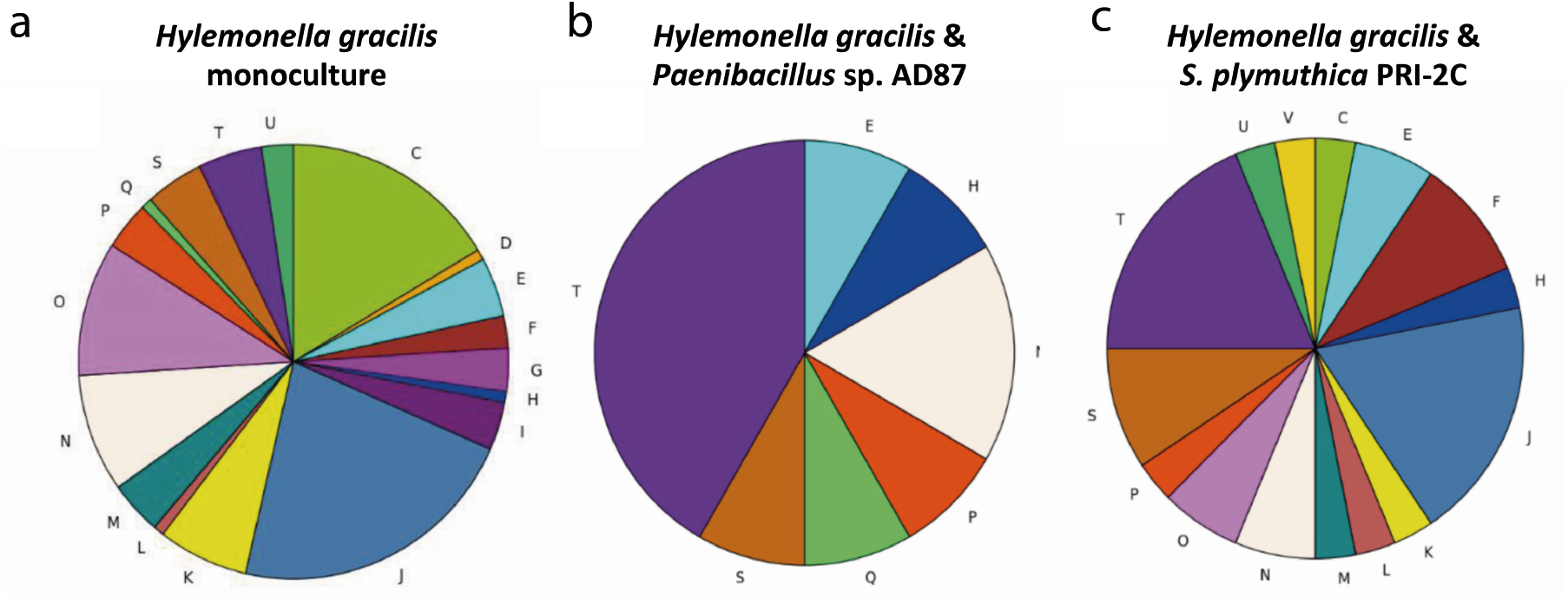
Pie-charts representing up-regulated genes identified by differential gene expression analysis and COG annotation. (**a**) *Hylemonella gracilis* monoculture gene expression level (**b**) *H. gracilis* in co-culture with *Paenibacillus* sp. AD87; (**c**) *H. gracilis* coculture with *S. plymuthica* PRI-2C. In the co-culture of *H. gracilis* with *Paenibacillus* sp. AD87, genes related to signal transduction (**T**) were the category with the most differentially expressed genes. In the co-culture of *H. gracilis* with *S. plymuthica* PRI-2C genes related to signal transduction (**T**), translation, ribosome structure and biogenesis (**J**) were the most prevalent differentially expressed gene categories. **COG-Abbreviations**: C: energy production and conversion; D: cell cycle control, cell division, chromosome partitioning; E: amino acid transport and metabolism; F: nucleotide transport and metabolism; G: carbohydrate transport and metabolism; H: coenzyme transport and metabolism; I: lipid transport and metabolism; J: translation, ribosomal structure and biogenesis; K: transcription; L: replication, recombination and repair; M: cell wall/membrane/envelope biogenesis; N: cell motility; NA: not assigned; O: posttranslational modification, protein turnover; chaperones; P: inorganic ion transport and metabolism; Q: secondary metabolites biosynthesis, transport and catabolism; R: general function prediction only; S: function unknown; T: signal transduction mechanisms; U: intracellular trafficking, secretion, and vesicular transport; V: defense mechanisms.

In *Paenibacillus* sp. AD87 histidine biosynthesis and dephosphorylation genes were up-regulated (**Supplementary Table 2**), while cellular-growth-related genes were down-regulated (**Supplementary Table 2**) at day 10 of the interaction with *H. gracilis* (**Fig. 6b**). For the interaction of *H. gracilis* with *Paenibacillus* sp. AD87 15 significant differentially expressed genes were found (0 at day five and 15 at day ten). At day five, genes related to sulfur assimilation, chemotaxis and response to (chemical/external) stimuli were upregulated in *H. gracilis* in the presence of *Paenibacillus* sp. AD87.

#### *Effect of inter-specific interactions on gene expression S. plymuthica* PRI-2C *and H. gracilis*

During the interaction of *S. plymuthica* PRI-2C with *H. gracilis*, 61 genes were significantly differentially expressed at day five and 10 at day ten. At day five, iron-sulfur clusterassembly-related genes, a sulfur transferase and a transaminase were up-regulated, while genes related to inorganic diphosphatase activity, exonuclease activity and DNA repair were downregulated. At day ten, genes related to sulfur transmembrane transport, sulfur compound catabolism and cysteine biosynthesis were upregulated, and genes related to sulfur compound metabolism and translation were downregulated. (**Supplementary Table 3** and **4**). For *S. plymuthica* PRI-2C, genes related to signal transduction and translation, ribosome structure and biogenesis were the most differentially expressed gene categories (**Fig. 6c**). For *H. gracilis* in interaction with *S. plymuthica* PRI-2C, 182 differentially expressed genes were identified at day ten and only one at day five. At day five, genes related to the ribosome/ribonucleoproteins, organelle organization/assembly and (iron)-sulfur cluster assembly were upregulated and genes related to the innate immune response (Toll Like Receptor signalling) were downregulated (**Supplementary Table 5** and **6**). At day ten, genes related to signal transduction and chemotaxis were upregulated in *H. gracilis*. For *H. gracilis*, the most upregulated genes were linked to chemotaxis pathway and iron scavenging, suggesting activity in competition (**Fig. 6a**).

### Metabolomic analysis of volatile compounds

The volatile blend composition of the monocultures differed from that of the co-cultures. Clear separations between the controls, monocultures and co-cultures were obtained in PLS-DA score plots (**Fig. 7a**). The analysis revealed a total of 25 volatile organic compounds produced by mono- and co-cultured bacteria that were not detected in the non-inoculated controls (**Table 3**). Of these, 17 were identified and categorized in six chemical classes (alkenes, benzoids, sulfides, thiocyanates, terpenes, furans). The remaining eight compounds could not be assigned with certainty to a known compound. The most abundant volatile organic compounds were sulfur-containing compounds such as dimethyl disulfide (C_2_H_6_S_3_) and dimethyl trisulfide (C_2_H_6_S_3_). These two sulfur compounds were produced by all three bacteria. Interestingly an unknown compound with a retention time (RT) of 26.4 min produced by the monocultures of *H. gracilis* was not detected in the interactions with *S. plymuthica* PRI-2C (**Table 3**). Two other unknown compounds with RT 4.15 min and with RT 24.34 min produced by the monocultures of *Paenibacillus* sp. AD87 were not detected in the co-cultivation with *H. gracilis* (**Table 3**).

**Figure 7:**
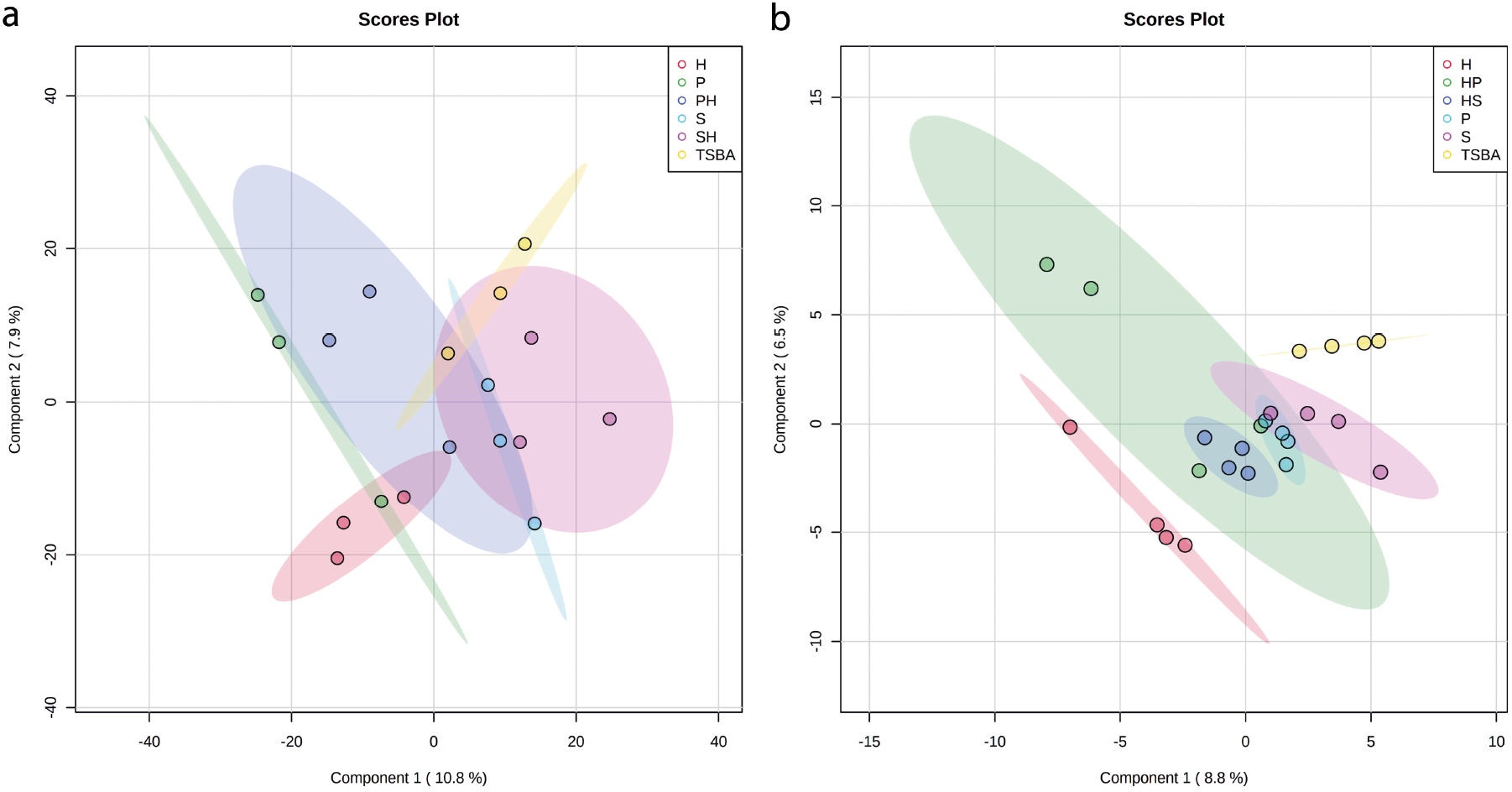
PLS-DA plots of the metabolomics data. (**a**) PLS-DA 2D-plots of volatiles emitted by monocultures and pairwise combinations of *H. gracilis, Paenibacillus* sp. AD87 and *Serratia plymuthica* after ten days of inoculation, time point (t=10 days) (**b**) PLS-DA 2D-plots of DART-MS data of monocultures and mixtures of *H. gracilis, Paenibacillus* sp. AD87 and *S. plymuthica* PRI-2C after ten days of inoculation, time point (t=10 days).

**Table 3:** Tentatively identified volatile organic compounds (VOCs) produced by a *H. gracilis, S. plymuthica* PRI-2C and *Paenibacillus* sp. AD87 strains in mono- and cocultures.

| # | Compound name | RT* | ELRI** | p-value*** | chemical class | Detected in bacterial culture |  |  |  |  |
| --- | --- | --- | --- | --- | --- | --- | --- | --- | --- | --- |
|  |  |  |  |  |  | H | S | P | SH | PH |
| 1 | 2-methylfuran | 2.18 | 738 | 0.041 | Furan | X |  | X | X | X |
| 2 | 2-methylpropanoic Acid | 3.01 | 755 | 0.014 | Alkenes | X | X | X | X | X |
| 3 | mix pentanal + heptane | 3.21 | 760 | 0.008 | Alkenes | X |  |  | X | X |
| 4 | methyl thycocyanate | 3.44 | 764 | 0.020 | Thioesters | X | X | X | X | X |
| 5 | 1-Pentanol | 3.95 | 772 | 0.012 | Alkenes |  | X | X | X | X |
| 6 | dimethyl disulfide | 4.01 | 775 | 0.012 | Sulfides | X | X | X | X | X |
| 7 | unknown compound 1 | 4.15 | 778 | 0.003 | - |  | X | X | X |  |
| 8 | toluene | 4.44 | 784 | 0.014 | Benzenoids | X | X | X | X | X |
| 9 | methyl Isovalerate | 4.76 | 789 | 0.018 | Terpenes |  | X | X | X | X |
| 10 | cyclohexane | 8.07 | 852 | 0.031 | Alkenes |  | X | X | X | X |
| 11 | dimethyl trisulfide | 11.35 | 914 | 0.013 | Sulfides | X | X | X | X | X |
| 12 | benzonitrile | 12.06 | 928 | 0.037 | Alkenes | X | X | X | X | X |
| 13 | 2-Ethyl-4-methylpentan-1-ol | 17.26 | 1026 | 0.015 | Alkenes |  | X | X | X | X |
| 14 | 2,5-bis(1-methylethyl)-pyrazine | 20.56 | 1090 | 0.031 | Pyrazines |  |  | X |  | X |
| 15 | undecane | 21.31 | 1100 | 0.014 | Alkenes | X |  | X | X | X |
| 16 | unknown compound 2 | 24.34 | 1140 | 0.013 | - |  |  | X |  |  |
| 17 | unknown compound 3 | 25.92 | 1160 | 0.011 | - | X | X | X | X | X |
| 19 | unknown compound 4 | 26.40 | 1165 | 0.018 | - | X |  | X | X | X |
| 20 | unknown compound 5 | 26.90 | 1170 | 0.003 | - | X |  | X |  | X |
| 21 | alpha-terpineol | 27.34 | 1178 | 0.016 | Terpenes | X | X | X | X | X |
| 22 | undecane, 2,6-dimethyl | 28.27 | 1190 | 0.004 | Benzenoids | X | X | X | X | X |
| 23 | gamma-terpineol | 28.42 | 1192 | 0.006 | Terpenes | X |  |  | X |  |
| 24 | terpene like compound 1 | 29.32 | 1202 | 0.012 | Terpenes | X |  |  | X |  |
| 25 | terpene like compound 2 | 31.49 | 1231 | 0.009 | Terpenes | X | X | X | X | X |
| Number of detected compounds (n) |  |  |  |  |  | 16 | 15 | 20 | 20 | 19 |

### DART-MS based metabolomics

Metabolomics analysis based on DART-MS revealed separations between the controls, monocultures, and co-cultures as presented in PLS-DA score plots (**Fig. 7b**). The metabolomic composition of the monocultures differed from that of the co-cultures (**Fig. 7b**). Statistical analysis (ONE-WAY ANOVA and post-hoc TUKEY HSD-test) revealed 617 significant mass features present on day five and day ten of which 48 could be tentatively assigned to specific compounds. Most of the significant peaks were found in the co-cultures of *H. gracilis* with *Paenibacillus* sp. AD87. The significant mass features and the corresponding tentative metabolites can be found in **Supplementary Table 10**.

### Mass spectrometry imaging metabolomics

LAESI-MSI was performed to visualize the localization of metabolites in their native environments in monoculture as well as during interaction without performing any extraction. Across all treatments, clear separation was observed amongst the samples for controls, monocultures and interactions **(Fig. 8a)**. An average of 1050 mass features was detected per treatment. To list mass features that could explain separation amongst the controls, monocultures and interactions, values of variable importance in projection (VIP) were calculated. The top 40 statistically significant mass features with VIP scores > 2.0 are shown in **Fig. 8b**. The box-and-whisker plots for the four statistically significant differentially abundant metabolites selected from the volcano plot for the pair HM and PH are shown in **Supplementary Figure 2a**. To visualize the statistically significant mass features between monocultures and co-cultures samples in a pairwise manner, volcano plots were constructed **(Supplementary Figure 3)**.

**Figure 8:**
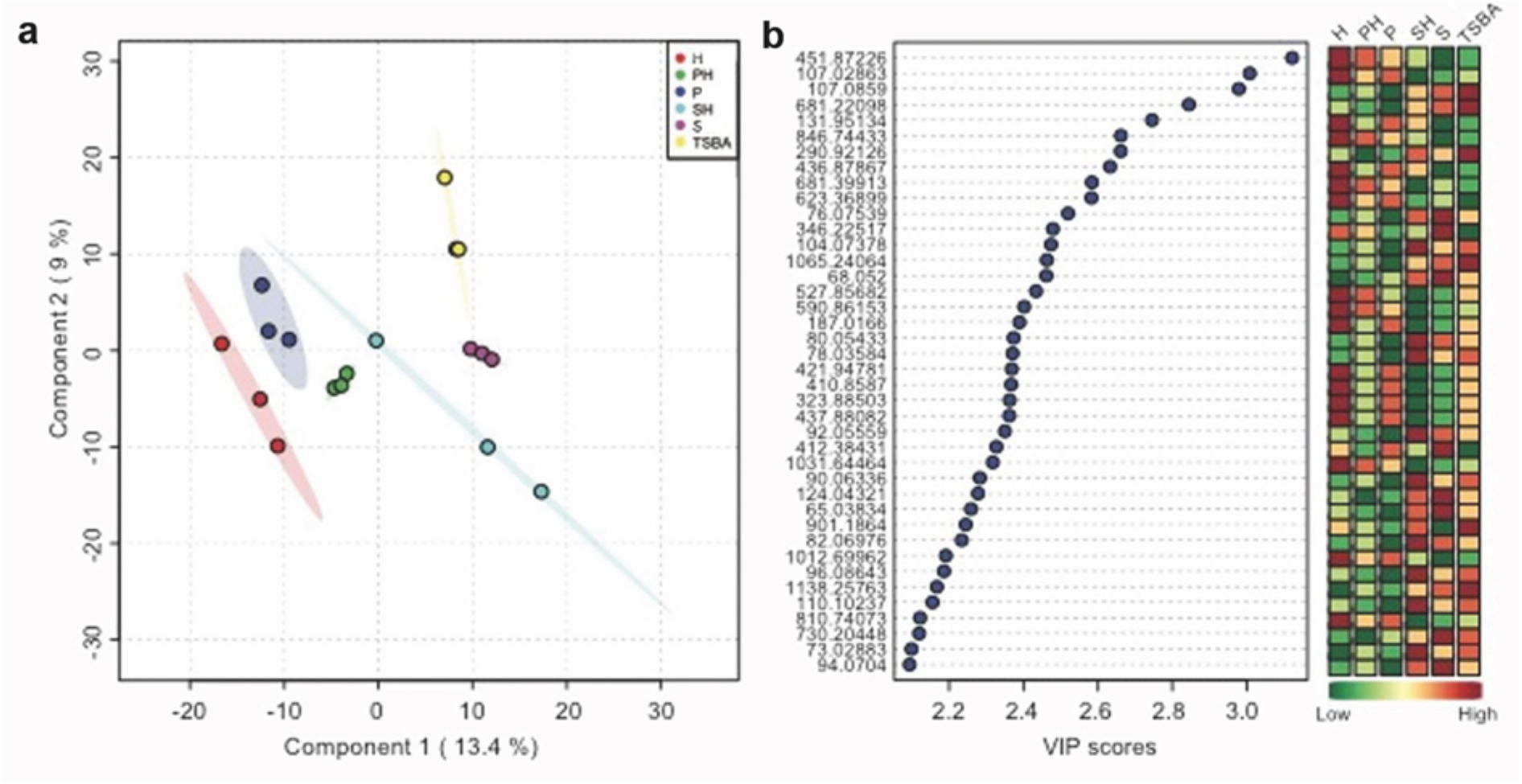
PLS-DA plots of the first 40 significant mass features observed in LAESI-MSI data. **(a)** PLS-DA score plot for *H. gracilis* monoculture (H), *Paenibacillus* sp. AD87 monoculture (P), *Paenibacillus* sp. AD87 – *H. gracilis* co-culture (PH), *S. plymuthica* PRI-2C monoculture (S), *S. plymuthica* PRI-2C – *H. gracilis* co-culture (SH) and TSBA control (TSBA). **(b)** Top 40 statistically significant features identified by PLS-DA based on Variable Importance in Projection (VIP) score. The colored boxes on the right indicate the relative concentrations of the corresponding metabolite in each group under study.

The volcano plot (**Supplementary Figure 3a**) for *H. gracilis* monoculture (HM) and the interaction of *H. gracilis* with *Paenibacillus* sp. AD87 (PH) shows 53 mass features (in green) located in the upper right quadrant, indicating that their concentrations are significantly higher in HM as compared to PH. 18 mass features (in red) in the upper left quadrant of the plot have a significantly lower concentration in HM as compared to PH. The ion intensity maps for these statistically significant metabolites are shown alongside box-and-whisker plots. The ion intensity maps are color coded based on the standard rainbow color scale where a pixel in red represented a high concentration and the pixel in black represents no concentration of the selected metabolite. As indicated, *m/z* 425.2886 and *m/z* 558.2832 show higher abundance in interaction sample PH, whereas *m/z* 410.8587 and *m/z* 716.7610 display high abundance in HM as compared to PH. For the pairwise analysis performed for *Paenibacillus* sp. AD87 monoculture (PM) and the co-culture of *H. gracilis* with *Paenibacillus* sp. AD87 (PH), 149 mass features (in green) displayed significantly high concentration in PM and 75 mass features (in red) had significantly low concentration in PM as compared to PH (**Supplementary Figure 3b**). This is also evident in the box-and-whisker plots and the ion intensity maps that are presented for four statistically significant metabolites belonging to this set (**Supplementary Figure 2b)**.

For the pairwise analysis for *H. gracilis* monoculture (HM) and the co-culture of *S. plymuthica* PRI-2C and *H. gracilis* (SH), 57 mass features (in green) displayed significantly high concentration in HM and 42 mass features had significantly low concentration in HM as compared to SH (**Supplementary Figure 3c**). The box-and-whisker plots along with the ion intensity maps for four statistically significant metabolites belonging to this set are shown in **Supplementary Figure 2c**. For the pairwise analysis for *S. plymuthica* PRI-2C monoculture (SM) and the interaction of *S. plymuthica* PRI-2C and *H. gracilis* (SH), 135 mass features (in green) displayed significantly high concentration in SM and 65 mass features had significantly low concentration in SM as compared to SH (**Supplementary Figure 3d**). The box-and-whisker plots along with the ion intensity maps for four statistically significant metabolites belonging to this set are shown in **Supplementary Figure 2d**.

To visualize the number of shared and unique metabolites amongst the monoculture and interaction samples Venn diagrams were plotted. The Venn diagram (**Supplementary Figure 3e**) for monocultures *H. gracilis* and *Paenibacillus* sp. AD87 and their interaction shows 80 metabolites unique to *H. gracilis* monoculture, 75 metabolites unique to *Paenibacillus* sp. AD87 monoculture and 100 metabolites that are unique during their interaction. 1062 metabolites were shared within these three treatments. Similarly, the Venn diagram **(Supplementary Figure 3f)** for monocultures *H. gracilis* and *S. plymuthica* PRI-2C and their interaction shows 196 metabolites unique to *H. gracilis* monoculture, 48 metabolites unique to *S. plymuthica* PRI-2C monoculture and 120 metabolites that are unique during their interaction.

## Discussion

Here we report the first time isolation of *H. gracilis* from a terrestrial soil sample. This bacterium passed a 0.1 μm filter, which suggests a very small cell size, theoretically justifying referring to these bacteria as ultra-small bacteria (26). However, against our expectation, the microscopical analysis revealed that this bacterium is not ultra-small in cell size but possesses a very thin diameter and showed the typical spiraled morphology known for these species (60–63). These observations are in line with previous research by Wang *et al*. showing that *H. gracilis* is capable of passing through filters of various pore sizes ranging from 0.45 μM to 0.1 μM (64), most probably thanks to their cell shape and cell morphology. *In silico* analysis of 16 terrestrial metagenome data available on MG-RAST (https://www.mg-rast.org/) showed that *H. gracilis* was not present in terrestrial metagenome data (not shown) suggesting that H. gracilis is not commonly present in terrestrial soils. The bacterial interaction assays revealed that *H. gracilis* grows faster when interacting with *Paenibacillus* sp. AD87 or *S. plymuthica* PRI-2C. The cell numbers of *H. gracilis* were higher when exposed to cell-free supernatants of *Paenibacillus* sp. AD87 and *S. plymuthica* PRI-2C, suggesting that the metabolites released by the latter bacteria in co-cultures with *H. gracilis* are associated with improved growth of *H. gracilis*. We hypothesized that *H. gracilis* grows better in co-culture, either because growth is stimulated by signals produced by the other organism, or because the environment that is created by the other organism allows *H. gracilis* to make more efficient use of certain metabolic pathways. Indeed, the metabolic experiments with BioLog™ plates showed that during interspecific interactions of *H. gracilis* with *Paenibacillus* sp. AD87 or with *S. plymuthica* PRI-2C, more carbohydrates could be utilized compared to the monocultures. This is an interesting observation, and it may indicate that interaction of bacteria can trigger the production of exo-enzymes enabling the degradation of carbohydrates, which the bacteria were not able to degrade in monoculture.

We speculated that since *H. gracilis* grows better in interaction with other bacteria and is of relatively small cell size, *H. gracilis* might have evolved according to a genome streamlining strategy, i.e. the adaptive loss of genes for which functions it relies on interaction with other bacteria in the immediate environment. The whole-genome sequencing of *H. gracilis* revealed a genome size of 3.82 Mbp. This is a relatively small genome size for free-living soil bacteria that typically have estimated average genome sizes of ~4.7 Mbp (34, 65–68). The *in silico* antiSMASH (43) comparison of genes that are part of secondary metabolite gene clusters showed that the *H. gracilis* genome contained only three gene clusters encoding the production of secondary metabolites (bacteriocins, terpenes, and aryl polyenes). Terpenes and aryl polyenes are known as protective compounds against abiotic stressors, while bacteriocins have antimicrobial activities against closely related bacteria (17, 69–73). We hypothesize that *H. gracilis* genome streamlining has allowed it to be more competitive, by retaining only the most essential metabolic functions while having roughly about one quarter less DNA to replicate during each cell division. Gene loss and reduced genome size may cause dependency on other microbes in their surroundings, and this may explain a considerable part of the phenomenon that most of the detectable bacteria in the environment are not cultivable under laboratory conditions.

To investigate the mechanisms of interaction, we performed transcriptome analysis on the interaction pairs of *H. gracilis* with *S. plymuthica* PRI-2C and *Paenibacillus* sp. AD87. Interestingly, a higher amount of significantly differentially expressed genes was induced by *H. gracilis* in the other two competing bacteria as compared to the transcriptomic changes in *H. gracilis*. Several processes, enriched according to GO term enrichment analysis, could be part of a mechanism(s) mediating interactions between *H. gracilis* and *S. plymuthica* PRI-2C and *Paenibacillus* sp. AD87, for example genes related to chemotaxis. Moreover, the GO terms for signal transduction, secondary metabolite production and, cell motility were enriched in the transcriptome of *H. gracilis* during the co-cultivation with *Paenibacillus* sp. AD87, suggesting that chemotaxis and cell movement is an important feature during interspecific interactions between these two bacterial taxa (74, 75). In addition, GO terms referring to Iron-sulfur (Fe-S) complex assembly were enriched in the transcriptomes of *H. gracilis* during the co-cultivation with *S. plymuthica* PRI-2C and *Paenibacillus* sp. AD87. FeS clusters are important for sustaining fundamental life processes: they participate in electron transfer, substrate binding/activation, iron or sulfur storage, regulation of gene expression, and enzyme activity (76, 77). This up-regulation could indicate that, potentially, in co-culture, normal-sized bacteria released metabolites that *H. gracilis* used for synthesizing Fe-S complexes. It is also possible that iron-sulfur complex assembly is activated during competition with the interacting bacteria for sulfur, or iron collection (scavenging) (78–81).

The metabolic pathway analysis showed that the loss of genes in *H. gracilis* does not appear to have resulted in functional loss of metabolic pathways. Loss of non-essential and possibly redundant genes in several metabolic pathways could explain why and how the genome of *H. gracilis* has become so small. The missing genes are not essential to complete metabolic pathways and only appear to result in limited options in certain metabolic pathways. RAST analysis showed that all basal metabolic pathways remain feasible with the annotated enzymes and pathways of *H. gracilis*. The only exception is EC term 5.2.1.1 (maleate isomerase) (it would help to specify in which pathway this enzyme is reported); There are several ways to synthesize fumarate, e.g. in the glycolysis pathway (63, 82, 83) and in the citric acid cycle (63, 84). Based on the available data, it cannot be unambiguously determined which alternative pathway may preferably be used by *H. gracilis* to synthesize fumarate.

The metabolomics analysis revealed the production of specific antimicrobial compounds such as pyrollnitrin (*S. plymuthica* PRI-2C) and 2,5-bis(1-methylethyl)-pyrazine (*Paenibacillus* sp. AD87) which are well known for their broad-spectrum antimicrobial activity (85–89). However, the produced antimicrobial compounds didn’t show activity against *H. gracilis*: in both interactions, *H. gracilis* showed increased growth when growing in co-culture with either *Paenibacillus* sp.AD87 or *S. plymuthica* PRI-2C.

The understanding of natural metabolites that mediate interactions between organisms in natural environments is the key to elucidate ecosystem functioning. The detection and identification of the compounds that mediate such interactions is still challenging. Techniques such as mass spectrometry imaging (MSI) provide new opportunities to study environmentally relevant metabolites in their spatial context (90–92). In this study, the metabolomics was performed using three independent approaches namely DART-MS analysis, GC/MS-Q-TOF analysis and Laser Ablation Electrospray Ionization-Mass Spectrometry Imaging mass spectrometry (LAESI-IMS) from living bacterial colonies. LAESI Imaging MS analysis revealed that several mass features were detected in higher abundance during the co-cultivation of *H. gracilis* with *Paenibacillus* sp. AD87, these mass features were m/z 425.2886 and m/z 558.2832. LAESI-MSI is not suitable for unambiguous compound annotation, but LAESI-MSI can still be used for putative compound annotation. To annotate the detected mass features to compounds with high certainty, LAESI mass spectrometry imaging should be coupled with ion mobility separation as suggested by (93–95). Yet, LAESI-MSI can help to spatially distinguish the produced secondary metabolites of living bacterial colonies with limited sample preparation and can give insight into the spatial distribution of metabolites.

Several studies indicate that the volatile blend composition of the volatiles greatly depends on biotic interactions and on growth conditions (15, 19, 96–98). Here, a higher number of volatile compounds were detected in the bacterial co-cultures, most likely due to the combination of emitted volatiles of the interacting bacteria. The high numbernumber of sulfur-containing compounds indicates that these compounds are commonly produced by bacteria and might play an important role in signaling during interspecific interactions (99, 100). No novel volatile compounds were detected during the co-culture of the three bacteria.

Overall, our study showed that *H. gracilis* is able to pass through 0.1 μM filter, and is present in terrestrial environments. The growth performance and physiological behavior of *H. gracilis* were dependent on the co-cultivated bacterial partner and they might be metabolically depending on the co-cultivated bacteria. At the same time, *H. gracilis* was able to change the physiology, release of volatile organic compounds and secreted enzymes of the co-cultivated bacteria without direct cell-cell contact.

Microbial interspecific interactions play an important role in the functioning of the terrestrial ecosystem. Soil microbial communities are very diverse and dynamic and involve frequent and sporadic interspecific interactions. Our study indicates that sulfur and Fe-S clusters could play important role in microbial interspecific interactions in terrestrial environments and more studies are required to understand their role.. The study of sporadic interspecific interactions and the inclusion of rare taxa in future analysis could help to better understand microbial communities and functions of those. Could you exemplify how this study improved our understanding? production of sulfur compounds and Fe-S clusters maybe?

## Data availability

The raw data of this article will be made available by the authors to any qualified researcher upon request. The whole genome sequence of *Hylemonella gracilis* strain NS1 is available at the NCBI GenBank under accession # CP031395, the raw reads of the transcriptomics data are available at the Sequence Read Archive (SRA) https://www.ncbi.nlm.nih.gov/sra under accession # PRJNA483535.

## Acknowledgement

This work was financially supported by The Netherlands Organization for Scientific Research (NWO) VIDI personal grant 864.11.015 granted to PG. The authors also want to thank the students of the WUR bioinformatics course BIF-51806 (2018) for their input and primary data analysis and discussions on the data.

## Author contributions

OT and PG designed the experiments. OT, AO, PK and WIJ performed the lab experiments. OT, PK and performed the data analysis and prepared the figures and tables. OT, PK, AO, VT, MHM, PB, KJFV and PG wrote the manuscript. All authors read and critically revised the manuscript.

