## Supplementary Material for "Exploring the interspecific interactions and the metabolome of the soil isolate *Hylemonella gracilis*"

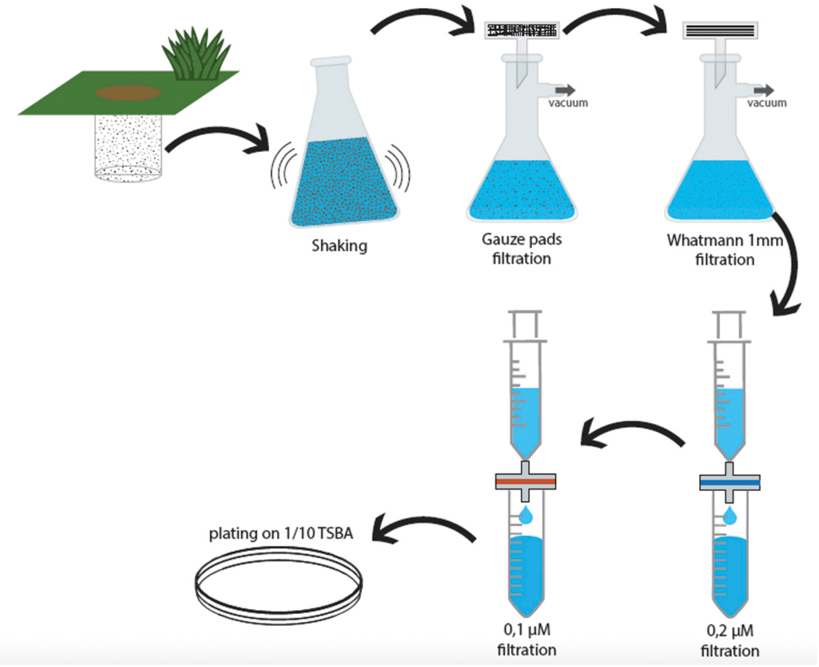


**Supplementary Figure 1:** Schematic overview of the applied isolation method used to isolate *H. gracilis* from soil.

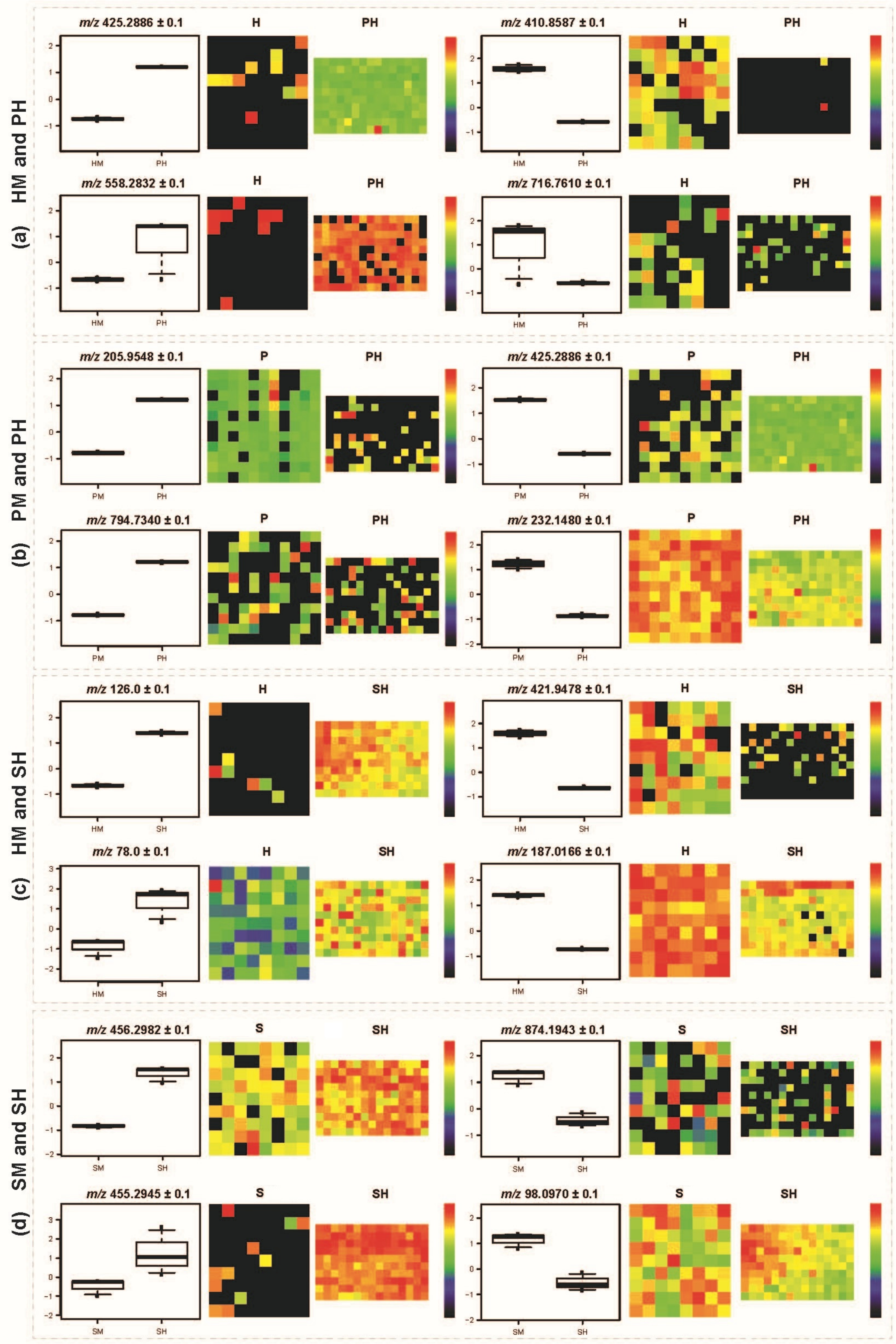


**Supplementary Figure 2:** Box plots for the significantly differentially abundant metabolites and their corresponding ion intensity maps found using Mass spectrometry imaging (MSI) during the interaction of *H. gracilis* with *Paenibacillus* sp. AD87 and *S. plymuthica* PRI-2C.

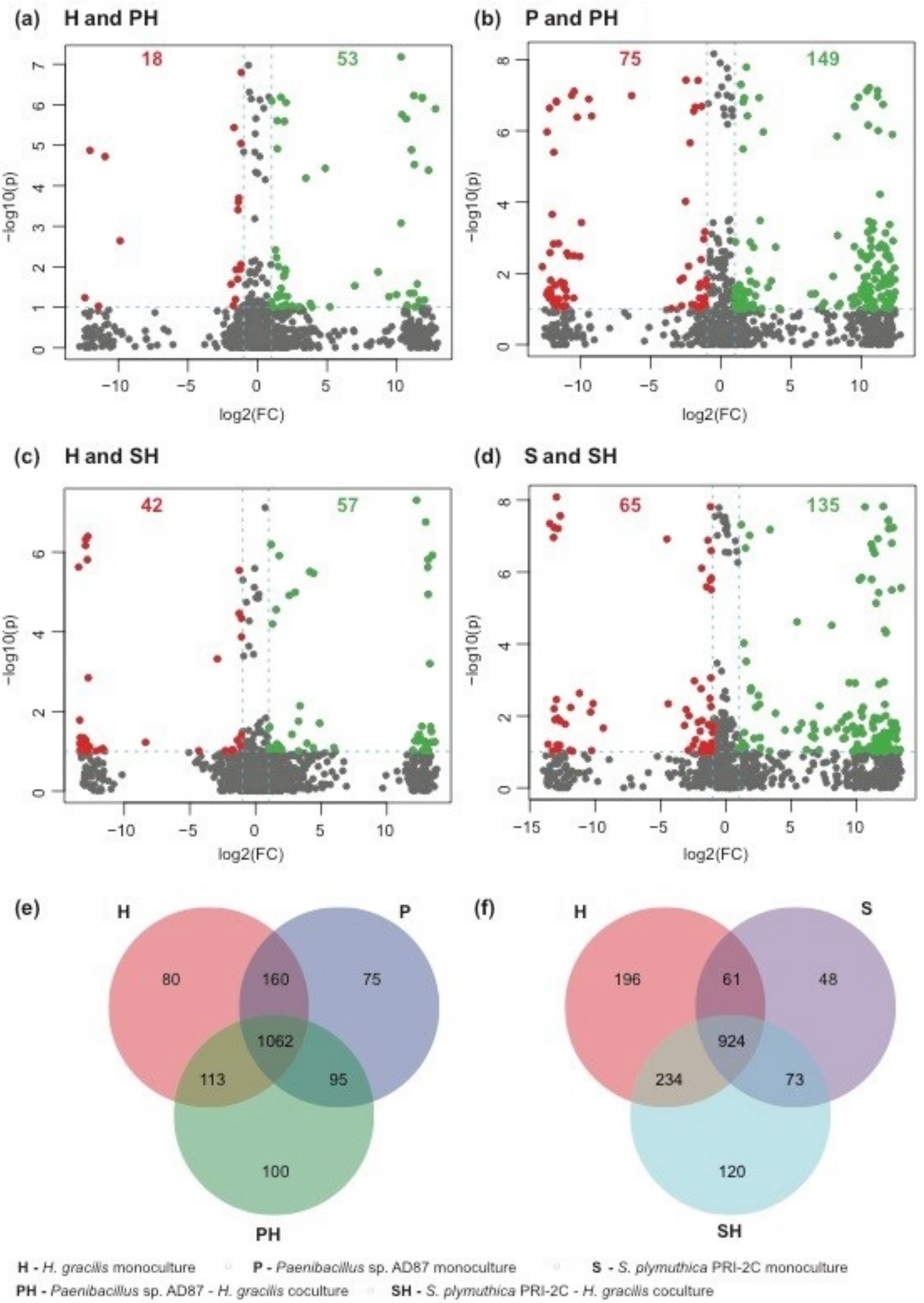


**Supplementary Figure 3: Volcano plots and Venn diagram to demonstrate metabolite concentration differences and unique/shared metabolites of the LAESI-MSI data**. **(a)** Volcano plot for *H. gracilis* monoculture (H) vs. *Paenibacillus* sp. AD87 - *H. gracilis* coculture (PH). **(b)** Volcano plot for *Paenibacillus* sp. AD87 monoculture (P) vs. *Paenibacillus* sp. AD87 - *H. gracilis* coculture (PH). **(c)** Volcano plot for *H. gracilis* monoculture (H) vs. *S. plymuthica* PRI-2C - *H. gracilis* coculture (SH). **(d)** Volcano plot for

*S. plymuthica* PRI-2C monoculture (S) vs. *S. plymuthica* PRI-2C - *H. gracilis* coculture (SH). Each point in the volcano plot represents one metabolite. Significantly differentially abundant metabolites were calculated with a fold change (FC) threshold of 2 on the x-axis and a t-tests threshold of 0.1 on the y-axis. The red and the green dots indicate statistically significant metabolites. The vertical FC threshold lines indicate an increase or decrease in concentration of metabolites. Negative log2 (FC) values indicated in red represent lower concentrations in native than in range expanding species; positive values indicated in green represent higher concentrations of metabolites in native than in range expanding species. **(e)** Venn diagram for HM-PM-PH. **(f)** Venn diagram for HM-SM-SH. To construct the Venn diagram, a single mass feature was considered even if it was present in only one replicate for a specific sample species.

**Supplementary Table 1:** Bacterial organisms used in this study.

| Strain / isolate / organism | Phylum/class | Genbank | Reference |
|---|---|---|---|
| *Hylemonella gracilis* isolate NS1 | beta-proteobacteria | CP0310395 | this study |
| *Paenibacillus* sp. AD87 | Firmicutes | KJ685299 | Tyc *et al.* 2014 |
| *Serratia plymuthica* PRI-2C | gamma-proteobacteria | CP015613 | Schmidt et al. 2017 |

**Supplementary Table 2**: Significantly differentially expressed genes of *Paenibacillus* sp. AD87, responding to *H. gracilis* at day 10.

| Gene | logFC | PValue | FDR | Function |
|---|---|---|---|---|
| gpAD87_RS06700 | -2.9328905 | 8.66E-15 | 6.48E-11 | rpoE; RNA polymerase sigma-70 factor, ECF subfamily |
| gpAD87_RS13150 | -1.8451399 | 7.78E-05 | 0.04477567 | opuBD; osmoprotectant transport system permease protein |
| gpAD87_RS13150 | -1.8451399 | 7.78E-05 | 0.04477567 | opuC; osmoprotectant transport system substrate-binding protein |
| gpAD87_RS30390 | -1.6913798 | 2.06E-06 | 0.0030772 | tatD; TatD DNase family protein [EC:3.1.21.-] |
| gpAD87_RS19945 | -1.2794042 | 1.17E-05 | 0.01253836 | UXS1; UDP-glucuronate decarboxylase [EC:4.1.1.35] |
| gpAD87_RS00275 | -1.2612217 | 1.63E-05 | 0.01628367 | E2.2.1.2; transaldolase [EC:2.2.1.2] |
| gpAD87_RS19920 | -1.2608789 | 1.78E-05 | 0.0166361 | pimB; phosphatidyl-myo-inositol dimannoside synthase [EC:2.4.1.346] |
| gpAD87_RS00270 | -1.2512648 | 5.28E-05 | 0.03593459 | PGD; 6-phosphogluconate dehydrogenase [EC:1.1.1.44 1.1.1.343] |
| gpAD87_RS19895 | -1.1972691 | 4.98E-06 | 0.00620873 | UGDH; UDPglucose 6-dehydrogenase [EC:1.1.1.22] |
| gpAD87_RS21205 | -1.1167672 | 2.65E-06 | 0.00360466 | N/A |
| gpAD87_RS26695 | -0.9979737 | 3.18E-05 | 0.02642266 | gerKA; spore germination protein KA |
| gpAD87_RS00725 | 1.0213585 | 9.92E-06 | 0.01142133 | E3.1.3.15B; histidinol-phosphatase (PHP family) [EC:3.1.3.15] |
| gpAD87_RS17335 | 1.43673471 | 9.05E-05 | 0.0483872 | abrB; transcriptional pleiotropic regulator of transition state genes |
| gpAD87_RS10500 | 1.48682555 | 6.63E-05 | 0.04132522 | deoC; deoxyribose-phosphate aldolase [EC:4.1.2.4] |
| gpAD87_RS11290 | 1.69953743 | 1.47E-06 | 0.00243757 | folD; methylenetetrahydrofolate dehydrogenase (NADP+) |

**Supplementary Table 3**: Significantly differentially expressed genes of *Serratia plymuthica* PRI-2C responding to *H. gracilis* at day 5.

| Gene | logFC | PValue | FDR | Function |
|---|---|---|---|---|
| Q5A_025180 | 1.484670421 | 4.37298312 | 1.16E-05 | rph; ribonuclease PH [EC:2.7.7.56] |

**Supplementary Table 4**: Significantly differentially expressed genes of *Serratia plymuthica* PRI-2C responding to *H. gracilis* at day 10.

| Gene | logFC | PValue | FDR | Function |
|---|---|---|---|---|
| Q5A_020525 | -3.539216745 | 3.79E-13 | 2.98E-10 | hcp; type VI secretion system secreted protein Hcp |
| Q5A_020555 | -2.840557251 | 6.03E-06 | 0.0011281 | fimA; major type 1 subunit fimbrin (pilin) |
| Q5A_023300 | -2.169738036 | 6.89E-08 | 2.10E-05 | malK; multiple sugar transport system ATP-binding protein [EC:3.6.3.-] |
| Q5A_011700 | -1.997102593 | 1.42E-05 | 0.0021876 | ynfM; MFS transporter, YNFM family, putative membrane transport protein |
| Q5A_003445 | -1.972950006 | 1.13E-06 | 0.00024804 | garD; galactarate dehydratase [EC:4.2.1.42] |
| Q5A_018430 | -1.744985992 | 9.02E-05 | 0.00993022 | N/A |
| Q5A_009750 | -1.734981082 | 4.74E-05 | 0.00622171 | ftnA; ferritin [EC:1.16.3.2] |
| Q5A_020535 | -1.729517104 | 0.000274679 | 0.02461418 | impB; type VI secretion system protein ImpB |
| Q5A_023295 | -1.625377127 | 2.34E-10 | 1.17E-07 | lamB; maltoporin |
| Q5A_003465 | -1.528329707 | 2.21E-05 | 0.00327673 | garL; 2-dehydro-3-deoxyglucarate aldolase [EC:4.1.2.20] |
| Q5A_003455 | -1.527498481 | 8.21E-05 | 0.00952465 | gudX; glucarate dehydratase-related protein |
| Q5A_003450 | -1.467669658 | 3.67E-06 | 0.00074239 | gudP; MFS transporter, ACS family, glucarate transporter |
| Q5A_010395 | -1.445444456 | 9.84E-06 | 0.00165079 | puuD; gamma-glutamyl-gamma-aminobutyrate hydrolase [EC:3.5.1.94] |
| Q5A_020530 | -1.425017256 | 5.82E-05 | 0.00719325 | impC; type VI secretion system protein ImpC |
| Q5A_002075 | -1.344637355 | 0.000219471 | 0.02052737 | ABC-2.P; ABC-2 type transport system permease protein |
| Q5A_015990 | -1.328859825 | 1.07E-07 | 3.15E-05 | paaC; ring-1,2-phenylacetyl-CoA epoxidase subunit PaaC [EC:1.14.13.149] |
| Q5A_010390 | -1.305376856 | 1.67E-07 | 4.46E-05 | puuR; HTH-type transcriptional regulator, repressor for puuD |
| Q5A_023310 | -1.296519929 | 5.59E-09 | 1.99E-06 | malE; maltose/maltodextrin transport system substrate-binding protein |
| Q5A_005185 | -1.29424411 | 1.11E-07 | 3.19E-05 | cybB; cytochrome b561 |
| Q5A_003460 | -1.287052853 | 0.000331618 | 0.02833289 | gudD; glucarate dehydratase [EC:4.2.1.40] |
| Q5A_015985 | -1.157880882 | 1.10E-05 | 0.00177766 | paaB; ring-1,2-phenylacetyl-CoA epoxidase subunit PaaB |
| Q5A_009735 | -1.129234286 | 2.54E-08 | 8.63E-06 | dksA; DnaK suppressor protein |
| Q5A_016000 | -1.118109907 | 1.34E-05 | 0.0021073 | paaE; ring-1,2-phenylacetyl-CoA epoxidase subunit PaaE |
| Q5A_015980 | -1.11728173 | 4.91E-06 | 0.00094221 | paaA; ring-1,2-phenylacetyl-CoA epoxidase subunit PaaA [EC:1.14.13.149] |
| Q5A_017330 | -1.103510732 | 4.63E-06 | 0.00091123 | katE; catalase [EC:1.11.1.6] |
| Q5A_015995 | -1.101590323 | 7.03E-06 | 0.00123738 | paaD; ring-1,2-phenylacetyl-CoA epoxidase subunit PaaD |
| Q5A_015975 | -1.093213554 | 3.94E-06 | 0.00078613 | paaZ; oxepin-CoA hydrolase / 3-oxo-5,6-dehydrosuberyl-CoA semialdehyde dehydrogenase |
| Q5A_016010 | -1.074845013 | 1.36E-05 | 0.00212107 | paaG; 2-(1,2-epoxy-1,2-dihydrophenyl)acetyl-CoA isomerase [EC:5.3.3.18] |
| Q5A_013825 | -1.003817338 | 0.000406824 | 0.03308758 | osmB; osmotically inducible lipoprotein OsmB |
| Q5A_008760 | -0.988582472 | 0.000254571 | 0.02322956 | hspQ; heat shock protein HspQ |
| Q5A_009380 | -0.977583963 | 0.000538345 | 0.04089512 | bssS; biofilm regulator BssS |
| Q5A_006035 | -0.974655768 | 1.64E-05 | 0.00249851 | acpD; FMN-dependent NADH-azoreductase [EC:1.7.-.-] |
| Q5A_003705 | -0.961603947 | 0.000667859 | 0.04823509 | csrA; carbon storage regulator |
| Q5A_005195 | -0.95850798 | 1.04E-05 | 0.00169051 | tomB; hha toxicity modulator TomB |
| Q5A_004910 | -0.924784476 | 1.74E-05 | 0.00262875 | panE; 2-dehydropantoate 2-reductase [EC:1.1.1.169] |
| Q5A_011805 | -0.908861788 | 0.000201997 | 0.01937745 | xdhB; xanthine dehydrogenase large subunit [EC:1.17.1.4] |
| Q5A_005190 | -0.887772818 | 0.00022171 | 0.02060805 | hha; haemolysin expression modulating protein |
| Q5A_015190 | -0.886911712 | 0.000277108 | 0.02468407 | phoH; phosphate starvation-inducible protein PhoH and related proteins |
| Q5A_013870 | -0.881053434 | 0.000295228 | 0.0259887 | ribA; GTP cyclohydrolase II [EC:3.5.4.25] |
| Q5A_010385 | -0.872345769 | 7.46E-05 | 0.00892893 | puuC; 4-(gamma-glutamylamino)butanal dehydrogenase [EC:1.2.1.99] |
| Q5A_024620 | -0.869220332 | 0.000360469 | 0.03030572 | membrane protein |
| Q5A_001015 | -0.867584758 | 0.000474421 | 0.03697766 | livM; branched-chain amino acid transport system permease protein |
| Q5A_023890 | -0.838466925 | 0.000467223 | 0.0367344 | PPIA; peptidyl-prolyl cis-trans isomerase A (cyclophilin A) [EC:5.2.1.8] |
| Q5A_022860 | -0.833565851 | 0.000526905 | 0.04023029 | yhcO; ribonuclease inhibitor |
| Q5A_017850 | -0.83112707 | 0.000165885 | 0.01633202 | mlaA; phospholipid-binding lipoprotein MlaA |
| Q5A_022140 | -0.824798608 | 0.000584632 | 0.04357256 | dkgA; 2,5-diketo-D-gluconate reductase A [EC:1.1.1.346] |
| Q5A_014735 | -0.785920924 | 0.000231103 | 0.02134478 | PTS-Man-EIIB; PTS system, mannose-specific IIB component [EC:2.7.1.191] |
| Q5A_014735 | -0.785920924 | 0.000231103 | 0.02134478 | PTS-Man-EIIA; PTS system, mannose-specific IIA component [EC:2.7.1.191] |
| Q5A_001825 | -0.784054272 | 0.000128357 | 0.01343315 | cysQ; 3(2) 5-bisphosphate nucleotidase [EC:3.1.3.7] |
| Q5A_010700 | -0.774422547 | 0.000383897 | 0.03209508 | htpX; heat shock protein HtpX [EC:3.4.24.-] |
| Q5A_017530 | -0.772978268 | 0.000651408 | 0.04732198 | yfbT; sugar-phosphatase [EC:3.1.3.23] |
| Q5A_014745 | -0.765542298 | 0.000232489 | 0.02134478 | PTS-Man-EIID; PTS system, mannose-specific IID component |
| Q5A_014680 | -0.744259112 | 0.000436668 | 0.03467316 | uncharacterized protein |
| Q5A_015210 | 0.739892297 | 0.000435304 | 0.03467316 | efeO; iron uptake system component EfeO |
| Q5A_007955 | 0.764324308 | 0.000306094 | 0.02663193 | manB; phosphomannomutase [EC:5.4.2.8] |
| Q5A_016985 | 0.76993376 | 0.000180387 | 0.0175292 | sbcB; exodeoxyribonuclease I [EC:3.1.11.1] |
| Q5A_019145 | 0.806510985 | 0.000131278 | 0.01364283 | pepB; PepB aminopeptidase [EC:3.4.11.23] |
| Q5A_024815 | 0.823927599 | 0.000110024 | 0.01193125 | argG; argininosuccinate synthase [EC:6.3.4.5] |
| Q5A_025450 | 0.865412879 | 0.000411826 | 0.0333134 | gidB; 16S rRNA (guanine527-N7)-methyltransferase [EC:2.1.1.170] |
| Q5A_013920 | 0.874038701 | 8.20E-05 | 0.00952465 | rluB; 23S rRNA pseudouridine2605 synthase [EC:5.4.99.22] |
| Q5A_000015 | 0.882396507 | 0.000271645 | 0.02457605 | recF; DNA replication and repair protein RecF |
| Q5A_019090 | 0.885515526 | 4.51E-05 | 0.00603012 | gcpE; (E)-4-hydroxy-3-methylbut-2-enyl-diphosphate synthase [EC:1.17.7.1 1.17.7.3] |
| Q5A_022525 | 0.919216585 | 0.000150986 | 0.01516709 | exuT; MFS transporter, ACS family, hexuronate transporter |
| Q5A_025475 | 0.919870586 | 0.000183702 | 0.01773611 | ATPF1D; F-type H+-transporting ATPase subunit delta |
| Q5A_023555 | 0.938796986 | 0.000428137 | 0.03444664 | RP-S4; small subunit ribosomal protein S4 |
| Q5A_023550 | 0.947685318 | 0.000356219 | 0.03011761 | rpoA; DNA-directed RNA polymerase subunit alpha [EC:2.7.7.6] |
| Q5A_007960 | 0.956964677 | 0.000620997 | 0.04568884 | ABC-2.LPSE.P; lipopolysaccharide transport system permease protein |
| Q5A_022100 | 0.958249224 | 0.000123936 | 0.01324783 | exbD; biopolymer transport protein ExbD |
| Q5A_005985 | 0.961851385 | 0.000159021 | 0.01578623 | rlpA; rare lipoprotein A |
| Q5A_003435 | 0.965989248 | 4.79E-05 | 0.00623402 | degP; serine protease Do [EC:3.4.21.107] |
| Q5A_023610 | 0.996820178 | 0.000401521 | 0.03283475 | RP-L24; large subunit ribosomal protein L24 |
| Q5A_003275 | 1.011595864 | 0.000138069 | 0.01424971 | murE; UDP-N-acetylmuramoyl-L-alanyl-D-glutamate--2,6-diaminopimelate ligase [EC:6.3.2.13] |
| Q5A_020635 | 1.02163108 | 0.000394468 | 0.03245928 | KARS; lysyl-tRNA synthetase, class II [EC:6.1.1.6] |
| Q5A_003765 | 1.031375231 | 0.000593738 | 0.0439866 | RP-S16; small subunit ribosomal protein S16 |
| Q5A_000705 | 1.032100444 | 0.000508445 | 0.0390199 | E2.7.7.24; glucose-1-phosphate thymidylyltransferase [EC:2.7.7.24] |
| Q5A_001315 | 1.073484442 | 3.28E-05 | 0.00462965 | rpoB; DNA-directed RNA polymerase subunit beta [EC:2.7.7.6] |
| Q5A_004935 | 1.076320393 | 0.000216005 | 0.02040276 | cyoC; cytochrome o ubiquinol oxidase subunit III |
| Q5A_004580 | 1.086598904 | 0.000114091 | 0.01228325 | uncharacterized protein |
| Q5A_019925 | 1.098903785 | 7.38E-05 | 0.00890562 | dnaE; DNA polymerase III subunit alpha [EC:2.7.7.7] |
| Q5A_010715 | 1.104205831 | 5.73E-05 | 0.00715045 | uncharacterized protein |
| Q5A_022645 | 1.109120361 | 2.59E-05 | 0.00372764 | gltD; glutamate synthase (NADPH/NADH) small chain [EC:1.4.1.13 1.4.1.14] |
| Q5A_005990 | 1.11082645 | 3.44E-05 | 0.00481475 | rodA; rod shape determining protein RodA |
| Q5A_025480 | 1.114497584 | 3.78E-05 | 0.00519142 | ATPF1A; F-type H+-transporting ATPase subunit alpha [EC:3.6.3.14] |
| Q5A_025495 | 1.115437397 | 9.85E-06 | 0.00165079 | ATPF1E; F-type H+-transporting ATPase subunit epsilon |
| Q5A_010095 | 1.123318289 | 0.000468845 | 0.0367344 | phoQ; two-component system, OmpR family, sensor histidine kinase PhoQ [EC:2.7.13.3] |
| Q5A_020770 | 1.124098577 | 8.92E-05 | 0.00993022 | gcvH; glycine cleavage system H protein |
| Q5A_020225 | 1.126066465 | 5.22E-05 | 0.00667858 | bisC; biotin/methionine sulfoxide reductase [EC:1.-.-.-] |
| Q5A_000715 | 1.14106216 | 6.48E-05 | 0.00794402 | wecE; dTDP-4-amino-4,6-dideoxygalactose transaminase [EC:2.6.1.59] |

| Gene | logFC | PValue | FDR | Function |
|---|---|---|---|---|
| Q5A_007965 | 1.159588453 | 7.48E-07 | 0.00018055 | ABC-2.LPSE.A; lipopolysaccharide transport system ATP-binding protein |
| Q5A_018270 | 1.163190653 | 0.00067399 | 0.04825964 | cysM; cysteine synthase B [EC:2.5.1.47] |
| Q5A_025490 | 1.181219213 | 6.49E-06 | 0.00115707 | ATPF1B; F-type H+-transporting ATPase subunit beta [EC:3.6.3.14] |
| Q5A_023600 | 1.189779741 | 0.000308508 | 0.02668686 | RP-S14; small subunit ribosomal protein S14 |
| Q5A_025485 | 1.231988595 | 2.69E-06 | 0.0005585 | ATPF1G; F-type H+-transporting ATPase subunit gamma |
| Q5A_021225 | 1.249899624 | 2.79E-06 | 0.00057138 | DLAT; pyruvate dehydrogenase E2 component (dihydrolipoamide acetyltransferase) [EC:2.3.1.12] |
| Q5A_007970 | 1.253396764 | 9.88E-06 | 0.00165079 | wbdD; O-antigen chain-terminating methyltransferase [EC:2.1.1.- 2.1.1.294 2.7.1.181] |
| Q5A_023595 | 1.255512256 | 0.000151012 | 0.01516709 | RP-S8; small subunit ribosomal protein S8 |
| Q5A_003510 | 1.269529949 | 8.75E-05 | 0.00991439 | ENO; enolase [EC:4.2.1.11] |
| Q5A_023665 | 1.279074138 | 0.000580067 | 0.04357256 | RP-L3; large subunit ribosomal protein L3 |
| Q5A_007565 | 1.311440579 | 0.000486067 | 0.03768909 | betB; betaine-aldehyde dehydrogenase [EC:1.2.1.8] |
| Q5A_010135 | 1.344256844 | 4.57E-05 | 0.00605029 | purB; adenylosuccinate lyase [EC:4.3.2.2] |
| Q5A_023590 | 1.394642505 | 6.27E-06 | 0.00114494 | RP-L6; large subunit ribosomal protein L6 |
| Q5A_013425 | 1.402843453 | 8.81E-05 | 0.00991439 | livK; branched-chain amino acid transport system substrate-binding protein |
| Q5A_013955 | 1.41352829 | 4.21E-05 | 0.0057213 | trpD; anthranilate phosphoribosyltransferase [EC:2.4.2.18] |
| Q5A_023150 | 1.413695908 | 0.000216775 | 0.02040276 | N/A |
| Q5A_005595 | 1.429913055 | 0.000502136 | 0.03873437 | N/A |
| Q5A_025240 | 1.448793539 | 0.000386643 | 0.03214509 | trmH; tRNA (guanosine-2-O-)-methyltransferase [EC:2.1.1.34] |
| Q5A_023585 | 1.450431734 | 5.46E-07 | 0.00013406 | RP-L18; large subunit ribosomal protein L18 |
| Q5A_013960 | 1.483167811 | 2.70E-09 | 1.01E-06 | trpCF; indole-3-glycerol phosphate synthase |
| Q5A_002020 | 1.489299833 | 7.87E-05 | 0.00927786 | truB; tRNA pseudouridine55 synthase [EC:5.4.99.25] |
| Q5A_001160 | 1.491198888 | 7.36E-10 | 3.15E-07 | rfaH; transcriptional antiterminator RfaH |
| Q5A_016980 | 1.494395971 | 0.000333217 | 0.02833289 | holE; DNA polymerase III subunit theta [EC:2.7.7.7] |
| Q5A_023575 | 1.520736432 | 8.56E-07 | 0.00020334 | RP-L15; large subunit ribosomal protein L15 |
| Q5A_023430 | 1.535747999 | 0.000323099 | 0.02778839 | aceB; malate synthase [EC:2.3.3.9] |
| Q5A_023660 | 1.545475067 | 0.00015085 | 0.01516709 | RP-L4; large subunit ribosomal protein L4 |
| Q5A_023580 | 1.598435296 | 1.08E-06 | 0.000246 | RP-S5; small subunit ribosomal protein S5 |
| Q5A_018150 | 1.616349566 | 8.70E-05 | 0.00991439 | fes; enterochelin esterase and related enzymes |
| Q5A_010010 | 1.633157137 | 2.07E-10 | 1.07E-07 | kdsA; 2-dehydro-3-deoxyphosphooctonate aldolase (KDO 8-P synthase) [EC:2.5.1.55] |
| Q5A_010365 | 1.661337332 | 0.000437904 | 0.03467316 | cspA; cold shock protein (beta-ribbon, CspA family) |
| Q5A_025245 | 1.673046601 | 0.000140408 | 0.01439184 | recG; ATP-dependent DNA helicase RecG [EC:3.6.4.12] |
| Q5A_023655 | 1.706448858 | 2.80E-05 | 0.00399534 | RP-L23; large subunit ribosomal protein L23 |
| Q5A_024965 | 1.722227382 | 9.71E-10 | 3.93E-07 | cysP; sulfate transport system substrate-binding protein |
| Q5A_008630 | 1.768955327 | 0.000585238 | 0.04357256 | ssuD; alkanesulfonate monooxygenase [EC:1.14.14.5] |
| Q5A_023630 | 1.798199582 | 2.19E-07 | 5.75E-05 | RP-L16; large subunit ribosomal protein L16 |
| Q5A_023650 | 1.81282525 | 8.88E-07 | 0.00020768 | RP-L2; large subunit ribosomal protein L2 |
| Q5A_007015 | 1.887269012 | 1.19E-05 | 0.00189254 | metQ; D-methionine transport system substrate-binding protein |
| Q5A_019155 | 1.923841527 | 5.19E-06 | 0.0009836 | fdx; ferredoxin, 2Fe-2S |
| Q5A_013965 | 1.936374402 | 4.44E-13 | 3.16E-10 | trpB; tryptophan synthase beta chain [EC:4.2.1.20] |
| Q5A_012540 | 1.952549176 | 8.72E-11 | 4.83E-08 | aldB; aldehyde dehydrogenase [EC:1.2.1.-] |
| Q5A_023625 | 1.955694281 | 3.46E-11 | 2.07E-08 | RP-L29; large subunit ribosomal protein L29 |
| Q5A_012535 | 1.958198587 | 1.22E-09 | 4.82E-07 | adhP; alcohol dehydrogenase, propanol-preferring [EC:1.1.1.1] |
| Q5A_025575 | 1.972560444 | 8.98E-05 | 0.00993022 | mnmE; tRNA modification GTPase [EC:3.6.-.-] |
| Q5A_018170 | 1.998960436 | 9.93E-06 | 0.00165079 | alsD; acetolactate decarboxylase [EC:4.1.1.5] |
| Q5A_023635 | 2.001100364 | 9.61E-10 | 3.93E-07 | RP-S3; small subunit ribosomal protein S3 |
| Q5A_013970 | 2.005143785 | 4.59E-07 | 0.00011445 | trpA; tryptophan synthase alpha chain [EC:4.2.1.20] |
| Q5A_002015 | 2.008984415 | 1.07E-10 | 5.71E-08 | rbfA; ribosome-binding factor A |
| Q5A_023645 | 2.073454609 | 5.03E-09 | 1.84E-06 | RP-S19; small subunit ribosomal protein S19 |
| Q5A_018290 | 2.078549829 | 9.21E-05 | 0.01005642 | cysP; sulfate transport system substrate-binding protein |
| Q5A_019160 | 2.097698613 | 3.77E-08 | 1.23E-05 | hscA; molecular chaperone HscA |
| Q5A_023640 | 2.114113863 | 4.69E-10 | 2.19E-07 | RP-L22; large subunit ribosomal protein L22 |
| Q5A_023620 | 2.11451368 | 7.20E-08 | 2.16E-05 | RP-S17; small subunit ribosomal protein S17 |
| Q5A_013945 | 2.171591652 | 2.32E-15 | 2.48E-12 | trpE; anthranilate synthase component I [EC:4.1.3.27] |
| Q5A_007560 | 2.223927239 | 1.05E-06 | 0.0002421 | betI; TetR/AcrR family transcriptional regulator, transcriptional repressor of bet genes |
| Q5A_023420 | 2.26789094 | 4.83E-06 | 0.00093786 | aceK; isocitrate dehydrogenase kinase/phosphatase [EC:2.7.11.5 3.1.3.-] |
| Q5A_023425 | 2.27591751 | 6.98E-10 | 3.07E-07 | E4.1.3.1; isocitrate lyase [EC:4.1.3.1] |
| Q5A_008095 | 2.28160194 | 3.26E-11 | 2.03E-08 | CTH; cystathionine gamma-lyase [EC:4.4.1.1] |
| Q5A_012240 | 2.29361659 | 2.90E-13 | 2.41E-10 | ABC.SS.S; simple sugar transport system substrate-binding protein |
| Q5A_007025 | 2.337387855 | 4.36E-05 | 0.00587802 | metN; D-methionine transport system ATP-binding protein |
| Q5A_006275 | 2.343443956 | 2.85E-07 | 7.22E-05 | kdpA; K+-transporting ATPase ATPase A chain [EC:3.6.3.12] |
| Q5A_025535 | 2.370161716 | 6.18E-06 | 0.00114135 | ABC.PA.S; polar amino acid transport system substrate-binding protein |
| Q5A_003565 | 2.383715546 | 1.80E-15 | 2.07E-12 | cysJ; sulfite reductase (NADPH) flavoprotein alpha-component [EC:1.8.1.2] |
| Q5A_008635 | 2.416532155 | 0.000159286 | 0.01578623 | ssuA; sulfonate transport system substrate-binding protein |
| Q5A_009885 | 2.468386668 | 1.27E-24 | 2.71E-21 | metQ; D-methionine transport system substrate-binding protein |
| Q5A_013150 | 2.488430101 | 0.000272611 | 0.02457605 | rbsB; ribose transport system substrate-binding protein |
| Q5A_005920 | 2.609600567 | 3.92E-12 | 2.67E-09 | yxeP; uncharacterized hydrolase [EC:3.-.-.-] |
| Q5A_003600 | 2.638693549 | 2.27E-09 | 8.72E-07 | cysC; adenylylsulfate kinase [EC:2.7.1.25] |
| Q5A_013950 | 2.684995402 | 3.42E-08 | 1.14E-05 | trpG; anthranilate synthase component II [EC:4.1.3.27] |
| Q5A_003585 | 2.693693766 | 2.94E-15 | 2.93E-12 | cysG; uroporphyrin-III C-methyltransferase |
| Q5A_008100 | 2.795399648 | 5.94E-21 | 8.08E-18 | CBS; cystathionine beta-synthase [EC:4.2.1.22] |
| Q5A_015155 | 2.962650632 | 6.70E-11 | 3.85E-08 | N/A |
| Q5A_018285 | 3.206409719 | 2.81E-16 | 3.50E-13 | cysU; sulfate transport system permease protein |
| Q5A_003570 | 3.27941167 | 4.09E-25 | 1.02E-21 | cysI; sulfite reductase (NADPH) hemoprotein beta-component [EC:1.8.1.2] |
| Q5A_023765 | 3.3435085 | 2.48E-07 | 6.41E-05 | tauC; taurine transport system permease protein |
| Q5A_003575 | 3.414713777 | 1.20E-21 | 1.80E-18 | cysH; phosphoadenosine phosphosulfate reductase [EC:1.8.4.8 1.8.4.10] |
| Q5A_013415 | 3.52258229 | 2.39E-05 | 0.00346779 | livG; branched-chain amino acid transport system ATP-binding protein |
| Q5A_003590 | 3.579275381 | 4.44E-27 | 1.33E-23 | cysD; sulfate adenylyltransferase subunit 2 [EC:2.7.7.4] |
| Q5A_012175 | 3.63206883 | 1.78E-05 | 0.00265913 | ABC.PA.A; polar amino acid transport system ATP-binding protein [EC:3.6.3.21] |
| Q5A_013405 | 3.660212984 | 2.24E-13 | 1.98E-10 | livH; branched-chain amino acid transport system permease protein |
| Q5A_018280 | 3.758245711 | 1.76E-23 | 3.29E-20 | cysW; sulfate transport system permease protein |
| Q5A_003595 | 3.86355858 | 5.63E-28 | 2.10E-24 | cysN; sulfate adenylyltransferase subunit 1 [EC:2.7.7.4] |
| Q5A_008640 | 3.97607918 | 1.13E-06 | 0.00024804 | ssuE; FMN reductase [EC:1.5.1.38] |
| Q5A_013420 | 4.062233442 | 1.23E-07 | 3.40E-05 | livF; branched-chain amino acid transport system ATP-binding protein |
| Q5A_012185 | 4.110365562 | 1.17E-07 | 3.30E-05 | ABC.PA.S; polar amino acid transport system substrate-binding protein |
| Q5A_023775 | 4.137817367 | 3.48E-33 | 1.74E-29 | tauA; taurine transport system substrate-binding protein |
| Q5A_015160 | 4.227876863 | 1.43E-39 | 2.14E-35 | cbl; LysR family transcriptional regulator, cys regulon transcriptional activator |
| Q5A_018275 | 4.32009885 | 7.32E-37 | 5.48E-33 | cysA; sulfate transport system ATP-binding protein [EC:3.6.3.25] |
| Q5A_012950 | 4.563975017 | 2.44E-10 | 1.18E-07 | cysE; serine O-acetyltransferase [EC:2.3.1.30] |
| Q5A_012180 | 4.668051343 | 1.31E-06 | 0.00028492 | ABC.PA.P; polar amino acid transport system permease protein |
| Q5A_013410 | 4.749568629 | 4.29E-08 | 1.36E-05 | livM; branched-chain amino acid transport system permease protein |
| Q5A_023760 | 5.264884391 | 9.51E-22 | 1.58E-18 | tauD; taurine dioxygenase [EC:1.14.11.17] |
| Q5A_023770 | 7.56664463 | 1.36E-13 | 1.27E-10 | tauB; taurine transport system ATP-binding protein [EC:3.6.3.36] |

**Supplementary Table 5**: Significantly differentially expressed genes of *H. gracilis* responding to *Serratia plymuthica* PRI-2C at day 5.

| Gene | logFC | PValue | FDR | Function |
|---|---|---|---|---|
| hylg_3594 | -10.30347602 | 1.79E-09 | 2.97E-06 | xylB; D-xylose 1-dehydrogenase [EC:1.1.1.175] |
| hylg_1896 | -9.05270584 | 3.82E-06 | 0.00173071 | livF; branched-chain amino acid transport system ATP-binding protein |
| hylg_681 | -8.967993981 | 1.91E-05 | 0.00681266 | rlmN; 23S rRNA (adenine2503-C2)-methyltransferase [EC:2.1.1.192] |
| hylg_2585 | -8.951345037 | 2.01E-05 | 0.00700548 | rbsC; ribose transport system permease protein |
| hylg_888 | -8.916187339 | 7.17E-06 | 0.00275048 | flhA; flagellar biosynthesis protein FlhA |
| hylg_2519 | -8.822014508 | 0.000219866 | 0.04112861 | ybeB; ribosome-associated protein |
| hylg_2782 | -8.738718593 | 8.00E-05 | 0.02064782 | uncharacterized protein |
| hylg_1702 | -8.701055125 | 0.000167787 | 0.03303859 | hisH; glutamine amidotransferase [EC:2.4.2.-] |
| hylg_3268 | -8.665105299 | 4.29E-05 | 0.01337529 | glnB; nitrogen regulatory protein P-II 1 |
| hylg_76 | -8.529781325 | 0.000125752 | 0.02727353 | NTE family protein |
| hylg_1992 | -8.287022835 | 0.000271079 | 0.0462024 | UGDH; UDPglucose 6-dehydrogenase [EC:1.1.1.22] |
| hylg_2781 | -4.886910881 | 1.08E-09 | 2.31E-06 | cheY; two-component system, chemotaxis family, chemotaxis protein CheY |
| hylg_3601 | -3.960678302 | 9.61E-05 | 0.02358015 | yiaN; TRAP-type transport system large permease protein |
| hylg_3527 | -3.848207509 | 0.000108449 | 0.02496838 | argG; argininosuccinate synthase [EC:6.3.4.5] |
| hylg_2468 | -3.624361126 | 6.29E-06 | 0.00254507 | btuB; vitamin B12 transporter |
| hylg_1855 | -3.563911838 | 0.00011898 | 0.02618433 | VARS; valyl-tRNA synthetase [EC:6.1.1.9] |
| hylg_3497 | -3.541208019 | 0.000152069 | 0.03117422 | N/A |
| hylg_54 | -3.530398733 | 0.000256743 | 0.04467628 | benE; benzoate membrane transport protein |
| hylg_1780 | -3.441042352 | 0.00013748 | 0.02939115 | polA; DNA polymerase I [EC:2.7.7.7] |
| hylg_2929 | -3.393004036 | 6.98E-05 | 0.01900203 | smc; chromosome segregation protein |
| hylg_2586 | -3.377821679 | 7.56E-05 | 0.02006357 | rbsA; ribose transport system ATP-binding protein [EC:3.6.3.17] |
| hylg_2197 | -3.328784638 | 2.33E-06 | 0.00108951 | hlpA; outer membrane protein |
| hylg_535 | -3.275554622 | 6.08E-05 | 0.01750013 | livK; branched-chain amino acid transport system substrate-binding protein |
| hylg_1719 | -3.254220622 | 4.84E-06 | 0.00201119 | sspA; stringent starvation protein A |
| hylg_538 | -2.949406851 | 0.000204119 | 0.03866636 | glnA; glutamine synthetase [EC:6.3.1.2] |
| hylg_3080 | -2.916611874 | 0.00024938 | 0.04393638 | sdhC; succinate dehydrogenase / fumarate reductase, cytochrome b subunit |
| hylg_1625 | -2.90876304 | 2.39E-05 | 0.00813567 | metK; S-adenosylmethionine synthetase [EC:2.5.1.6] |
| hylg_1958 | -2.845623805 | 8.52E-05 | 0.0216103 | fimV; pilus assembly protein FimV |
| hylg_2584 | -2.8004223 | 0.000291159 | 0.04788124 | rbsB; ribose transport system substrate-binding protein |
| hylg_3274 | -2.715164625 | 0.000162179 | 0.03236019 | ppa; inorganic pyrophosphatase [EC:3.6.1.1] |
| hylg_1357 | -2.646079415 | 0.000249555 | 0.04393638 | N/A |
| hylg_1747 | -2.24362962 | 0.000145461 | 0.03023366 | N/A |
| hylg_1116 | -2.149397607 | 0.000230074 | 0.04250696 | E3.3.1.1; adenosylhomocysteinase [EC:3.3.1.1] |
| hylg_3616 | -2.133709269 | 0.000238398 | 0.04315128 | N/A |
| hylg_197 | -2.014259372 | 6.94E-06 | 0.00273302 | coxB; cytochrome c oxidase subunit II [EC:1.9.3.1] |
| hylg_3051 | -1.961997657 | 3.78E-05 | 0.01229204 | ENO; enolase [EC:4.2.1.11] |
| hylg_2411 | 1.836427686 | 0.000271688 | 0.0462024 | iscS; cysteine desulfurase [EC:2.8.1.7] |
| hylg_2413 | 2.142314215 | 0.000103623 | 0.02461468 | iscA; iron-sulfur cluster assembly protein |
| hylg_956 | 2.252713163 | 6.32E-05 | 0.01784859 | N/A |
| hylg_3349 | 2.513218505 | 1.03E-05 | 0.00387033 | RP-S21; small subunit ribosomal protein S21 |
| hylg_244 | 2.645172803 | 1.82E-06 | 0.00090883 | rluF; 23S rRNA pseudouridine2604 synthase [EC:5.4.99.21] |
| hylg_3064 | 2.836010247 | 4.50E-05 | 0.01374931 | N/A |
| hylg_764 | 2.845097872 | 1.69E-06 | 0.00086973 | ksgA; 16S rRNA (adenine1518-N6/adenine1519-N6)-dimethyltransferase [EC:2.1.1.182] |
| hylg_2414 | 2.990210312 | 9.27E-05 | 0.02312815 | hscB; molecular chaperone HscB |
| hylg_3350 | 3.165994836 | 2.98E-07 | 0.00017819 | uncharacterized protein |
| hylg_3472 | 3.309925768 | 7.35E-07 | 0.00042277 | bpeF; multidrug efflux pump |
| hylg_2415 | 3.400742126 | 1.22E-06 | 0.0006771 | hscA; molecular chaperone HscA |
| hylg_3352 | 3.581267445 | 4.51E-08 | 4.22E-05 | gudD; glucarate dehydratase [EC:4.2.1.40] |
| hylg_359 | 3.815734971 | 1.02E-07 | 7.36E-05 | cheW; purine-binding chemotaxis protein CheW |
| hylg_3471 | 3.830015991 | 3.18E-05 | 0.01057743 | bpeE; membrane fusion protein, multidrug efflux system |
| hylg_3068 | 3.969695582 | 4.42E-06 | 0.00189132 | trmD; tRNA (guanine37-N1)-methyltransferase [EC:2.1.1.228] |
| hylg_1092 | 5.118307349 | 1.69E-09 | 2.97E-06 | N/A |
| hylg_360 | 5.190346574 | 5.26E-05 | 0.01543046 | mcp; methyl-accepting chemotaxis protein |
| hylg_1563 | 5.297224632 | 1.03E-07 | 7.36E-05 | mcp; methyl-accepting chemotaxis protein |
| hylg_3477 | 5.333738341 | 1.97E-06 | 0.00094974 | acrA; membrane fusion protein, multidrug efflux system |
| hylg_2376 | 5.556284733 | 2.87E-09 | 4.29E-06 | N/A |
| hylg_357 | 5.920231993 | 7.62E-11 | 2.28E-07 | N/A |
| hylg_363 | 5.987661764 | 1.51E-12 | 7.53E-09 | mcp; methyl-accepting chemotaxis protein |
| hylg_1561 | 6.333104946 | 4.06E-10 | 1.01E-06 | N/A |
| hylg_358 | 6.400072284 | 3.67E-13 | 2.75E-09 | N/A |
| hylg_361 | 7.03578819 | 7.95E-09 | 1.02E-05 | cheW; purine-binding chemotaxis protein CheW |

**Supplementary Table 6**: Significantly up- or down regulated genes of *H. gracilis* responding to *Serratia plymuthica* PRI-2C at day 10.

| Gene | logFC | PValue | FDR | Function |
|---|---|---|---|---|
| hylg_1092 | 4.028021412 | 5.89E-07 | 0.00152989 | N/A |
| hylg_1561 | 4.825700429 | 3.64E-06 | 0.00680041 | N/A |
| hylg_1562 | 4.569714447 | 8.80E-07 | 0.00188109 | cheW; purine-binding chemotaxis protein CheW |
| hylg_1563 | 5.426932759 | 7.79E-08 | 0.00029152 | mcp; methyl-accepting chemotaxis protein |
| hylg_2376 | 3.649019855 | 1.95E-05 | 0.02289822 | N/A |
| hylg_357 | 4.707340509 | 6.13E-07 | 0.00152989 | N/A |
| hylg_358 | 5.714802483 | 2.38E-09 | 1.78E-05 | N/A |
| hylg_359 | 3.924767967 | 9.74E-06 | 0.01458213 | cheW; purine-binding chemotaxis protein CheW |
| hylg_361 | 6.080192446 | 1.41E-05 | 0.01917412 | cheW; purine-binding chemotaxis protein CheW |
| hylg_363 | 5.259687546 | 4.19E-10 | 6.27E-06 | mcp; methyl-accepting chemotaxis protein |

**Supplementary Table 7:** Significantly up- or down regulated genes of *H. gracilis* responding to *Paenibacillus* sp. AD87 at day 5.

| Gene | logFC | PValue | FDR | Function |
|---|---|---|---|---|
| hylg_283 | 1.83868209 | 1.21E-05 | 0.02840838 | soxY; sulfur-oxidizing protein SoxY |
| hylg_2715 | 3.18297333 | 3.22E-08 | 0.000482259 | pobA; p-hydroxybenzoate 3-monooxygenase [EC:1.14.13.2] |
| hylg_1092 | 3.66916806 | 3.70E-06 | 0.013834781 | |
| hylg_1561 | 3.88090845 | 2.32E-05 | 0.043402807 | |
| hylg_689 | 3.90578955 | 5.43E-07 | 0.002708319 | |
| hylg_1563 | 3.91597386 | 2.64E-05 | 0.043946331 | mcp; methyl-accepting chemotaxis protein |
| hylg_361 | 4.72545337 | 1.33E-05 | 0.02840838 | cheW; purine-binding chemotaxis protein CheW |
| hylg_2376 | 4.8197097 | 9.29E-08 | 0.000694792 | |

**Supplementary Table 8:** Significantly up- or down regulated genes of *Paenibacillus* responding to *H. gracilis* at day 10.

| Gene | logFC | PValue | FDR | Function |
|---|---|---|---|---|
| gpAD87_RS06700 | -2.9328905 | 8.66E-15 | 6.48E-11 | rpoE; RNA polymerase sigma-70 factor, ECF subfamily |
| gpAD87_RS13150 | -1.8451399 | 7.78E-05 | 0.04477567 | opuBD; osmoprotectant transport system permease protein |
| gpAD87_RS13150 | -1.8451399 | 7.78E-05 | 0.04477567 | opuC; osmoprotectant transport system substrate-binding protein |
| gpAD87_RS30390 | -1.6913798 | 2.06E-06 | 0.0030772 | tatD; TatD DNase family protein [EC:3.1.21.-] |
| gpAD87_RS19945 | -1.2794042 | 1.17E-05 | 0.01253836 | UXS1; UDP-glucuronate decarboxylase [EC:4.1.1.35] |
| gpAD87_RS00275 | -1.2612217 | 1.63E-05 | 0.01628367 | E2.2.1.2; transaldolase [EC:2.2.1.2] |
| gpAD87_RS19920 | -1.2608789 | 1.78E-05 | 0.0166361 | pimB; phosphatidyl-myo-inositol dimannoside synthase [EC:2.4.1.346] |
| gpAD87_RS00270 | -1.2512648 | 5.28E-05 | 0.03593459 | PGD; 6-phosphogluconate dehydrogenase [EC:1.1.1.44 1.1.1.343] |
| gpAD87_RS19895 | -1.1972691 | 4.98E-06 | 0.00620873 | UGDH; UDPglucose 6-dehydrogenase [EC:1.1.1.22] |
| gpAD87_RS21205 | -1.1167672 | 2.65E-06 | 0.00360466 | N/A |
| gpAD87_RS26695 | -0.9979737 | 3.18E-05 | 0.02642266 | gerKA; spore germination protein KA |
| gpAD87_RS00725 | 1.0213585 | 9.92E-06 | 0.01142133 | E3.1.3.15B; histidinol-phosphatase (PHP family) [EC:3.1.3.15] |
| gpAD87_RS17335 | 1.43673471 | 9.05E-05 | 0.0483872 | abrB; transcriptional pleiotropic regulator of transition state genes |
| gpAD87_RS10500 | 1.48682555 | 6.63E-05 | 0.04132522 | deoC; deoxyribose-phosphate aldolase [EC:4.1.2.4] |
| gpAD87_RS11290 | 1.69953743 | 1.47E-06 | 0.00243757 | folD; methylenetetrahydrofolate dehydrogenase (NADP+) |

**Supplementary Table 9:** Tentatively identified metabolites revealed by DART-MS analysis.

| Compound # | M/Z | RT | Compound | p.value | FDR-value |
|---|---|---|---|---|---|
| 1 | 165.1384 | 0.20 | Actinidine | 0.000 | 0.000 |
| 2 | 223.0961 | 0.30 | Apiole | 0.007 | 0.037 |
| 3 | 249.148 | 0.40 | 1,2-Dihydrosantonin | 0.007 | 0.037 |
| 4 | 137.1072 | 0.60 | N,N-Dimethyl-1,4-phenylenediamine | 0.006 | 0.037 |
| 5 | 110.0602 | 0.60 | N-Vinyl-2-pyrrolidone | 0.007 | 0.037 |
| 6 | 212.2005 | 0.70 | Oxidized Latia luciferin | 0.007 | 0.037 |
| 7 | 161.0806 | 0.70 | Pimelate | 0.000 | 0.000 |
| 8 | 132.1018 | 0.80 | 2-Hydroxycyclohexan-1-one | 0.000 | 0.000 |
| 9 | 186.1486 | 0.80 | 6-Oxocineole | 0.000 | 0.000 |
| 10 | 172.1329 | 0.80 | Boschnialactone | 0.006 | 0.037 |
| 11 | 227.0133 | 0.90 | Bismuth | 0.000 | 0.000 |
| 12 | 102.0916 | 0.90 | Cyclopentanone | 0.000 | 0.000 |
| 13 | 118.0863 | 1.00 | 5-Valerolactone | 0.000 | 0.000 |
| 14 | 178.1337 | 1.10 | Tryptamine | 0.006 | 0.037 |
| 15 | 152.1432 | 1.20 | p-Cymene | 0.011 | 0.042 |
| 16 | 87.0087 | 1.30 | Pyruvate | 0.000 | 0.000 |
| 17 | 197.0222 | 1.50 | 5-Methyl-3-isoxazolyl sulfate | 0.009 | 0.037 |
| 18 | 175.0247 | 1.50 | Ascorbate | 0.006 | 0.037 |
| 19 | 121.0294 | 1.50 | Benzoate | 0.007 | 0.037 |
| 20 | 179.0559 | 1.50 | D-Glucose | 0.014 | 0.048 |
| 21 | 255.2328 | 1.50 | Hexadecanoic acid | 0.007 | 0.037 |
| 22 | 100.0759 | 1.50 | Pentanamide | 0.000 | 0.000 |
| 23 | 304.2477 | 1.60 | 2,3-Dihydroxycyclopentaneundecanoic | 0.000 | 0.000 |
| 24 | 217.1795 | 1.70 | 12-Hydroxydodecanoic acid | 0.000 | 0.000 |
| 25 | 123.0553 | 1.80 | Nicotinamide | 0.006 | 0.037 |
| 26 | 166.086 | 1.80 | trans-Cinnamate | 0.006 | 0.037 |
| 27 | 131.0713 | 1.90 | R-2-Hydroxyisocaproate | 0.000 | 0.000 |
| 28 | 129.0557 | 2.00 | 4-Methyl-2-oxopentanoate | 0.006 | 0.037 |
| 29 | 253.2171 | 2.00 | 9Z-Hexadecenoic acid | 0.006 | 0.037 |
| 30 | 164.0715 | 2.00 | Coumarin | 0.000 | 0.000 |
| 31 | 171.139 | 2.00 | Decanoic acid | 0.014 | 0.047 |
| 32 | 130.0873 | 2.30 | L-Leucine | 0.000 | 0.001 |
| 33 | 165.0191 | 2.40 | Phthalate | 0.000 | 0.000 |
| 34 | 117.0557 | 2.50 | 5-Hydroxypentanoate | 0.000 | 0.000 |
| 35 | 101.0607 | 2.50 | Pentanoate | 0.000 | 0.000 |
| 36 | 134.0471 | 2.60 | Adenine | 0.000 | 0.000 |
| 37 | 241.2172 | 2.60 | Pentadecanoic acid | 0.000 | 0.000 |
| 38 | 128.0353 | 2.70 | 4-Oxoproline | 0.000 | 0.005 |
| 39 | 130.0873 | 2.70 | Cyclohexane-1,3-dione | 0.000 | 0.000 |
| 40 | 227.2015 | 2.70 | Tetradecanoic acid | 0.007 | 0.037 |
| 41 | 116.9285 | 2.70 | unknown | 0.000 | 0.000 |
| 42 | 187.1338 | 3.30 | 10-Hydroxydecanoic acid | 0.000 | 0.000 |
| 43 | 243.1962 | 3.30 | 2S-Hydroxytetradecanoic acid | 0.000 | 0.000 |
| 44 | 87.0329 | 3.40 | Hydroxylamine hydrochloride | 0.000 | 0.000 |
| 45 | 89.0243 | 3.50 | Glycerone | 0.000 | 0.000 |
| 46 | 116.0716 | 3.50 | L-Valine | 0.006 | 0.037 |
| 47 | 224.999 | 3.60 | Aminopyrrolnitrin | 0.000 | 0.000 |
| 48 | 254.9731 | 3.60 | Pyrrolnitrin | 0.000 | 0.000 |

### Supplementary Methods

#### DNA isolation and genome sequencing of *H. gracilis*

For genomic DNA extraction a single colony of *H. gracilis* was picked from plate and inoculated in triplicates in 25 mL LB medium and incubated at 24 °C, 200 rpm.. Genomic DNA was extracted from three days grown cultures by using the QIAGEN Genomic-tip 500/G DNA extraction kit (Qiagen, cat# 10262). The extracted DNA was dissolved in 200 µl sterile nuclease-free water and quantified with a NanoDrop Spectrophotometer (ND 2000, Thermo Fisher Scientific, The Netherlands). Additionally, a 1.0% TBE agarose gel was run to check the size and integrity of the isolated genomic DNA. The extracted genomic DNA was stored at -20 °C and subjected to PacBio real-time DNA sequencing using the PacBio RS II sequencing platform at the Institute for Genome Sciences (IGS), Baltimore, Maryland, USA.

#### Pathway annotations

For pathway annotations, BlastKOALA was used (https://www.kegg.jp/blastkoala/) (8). FASTA files of the Protein sequences of the significantly differentially expressed genes were extracted from GenBank (https://www.ncbi.nlm.nih.gov/genbank/) (9) and merged into a new FASTA file corresponding to each comparison group. The FASTA files were uploaded to BlastKOALA and a search was performed against the prokaryotes KEGG GENES database file.

#### GC-Q-TOF data analysis

Volatile organic compounds (VOCs) were trapped by using glass Petri dishes (10). At the top of the Petri dishes the lid was connected with an outlet to a steel trap containing 150 mg Tenax TA and 150 mg Carbopack B (Markes International Ltd). The Tenax steel traps were added after four and nine days of incubation and collected at day five and ten of incubation. As controls glass Petri dishes containing 1/10$^{th}$ TSBA without bacteria were used. The VOCs were desorbed from the Tenax using an automated thermos desorption unit (Unity TD-100, Markes International Ltd., Llantrisant, UK). The desorbed volatiles were introduced into the GC-QTOF (model Agilent 7890B GC and the Agilent 7200A QTOF, Santa Clara, USA) and measured as described in (11). For VOC data analysis and compound identification Mass-spectra were extracted with MassHunter Qualitative Analysis Software V B.06.00 Build 6.0.633.0 (Agilent Technologies, Santa Clara, USA) and exported as netCDF files using the MassHunter GC/MS (GC-AIA) Translator B.07.00 SP2 163.. The netCDF files were imported to MZmine V2.24 (Copyright © 2005-2015 MZmine Development Team) (12) and compounds were identified based on their mass spectra and by their linear retention indexes (LRI) in combination with two mass-spectral-libraries: NIST 2014 V2.20 (National Institute of Standards and Technology, USA http://www.nist.gov). The LRI values were calculated using AMDIS 2.72 (National Institute of Standards and Technology, USA). Peak lists containing the mass features of each treatment were exported in csv file format and uploaded to Metaboanalyst V3.0 (www.metaboanalyst.ca) running on a local UNIX server for statistical analysis (13). To identify significantly differentially (OK?) abundant masses ONE-WAY-ANOVA with post-hoc TUKEY test was performed between the data sets.

**Ambient mass-spectrometry imaging analysis LEASI-MS data analysis**

The positive ions were acquired in a mass range of 50 to 1200 m/z. The MS data was lock mass corrected post data acquisition using leucine encephalin (C25H37N5O7m/z= 556.2771), which was used as an internal standard. All the acquired Waters *.RAW data files were converted to open file format *.imzML using an in-house script written in R. Later, this data was pre-processed in multiple steps to remove noise and to make the data comparable. First, square root transformation was applied to the data to stabilize the variance. Then, baseline correction was performed to enhance the contrast of peaks to the baseline. For better comparison of intensity values and to remove small batch effects, Total-Ion-Current (TIC)-based normalization was applied. This was followed by spectral alignment and peak detection to extract a list of significant mass features for each sample replicate per treatment. In the end, a mass feature matrix was generated with sample replicates for each treatment in columns and mass features in rows. This feature matrix was used to perform further statistical analysis. The pre-processing and peak-detection steps were applied using R scripts developed in-house and the functions available within the MALDIquant R package (62). To perform multivariate analysis, the feature matrix was imported into the online version of Metaboanalyst 4.0 (63). Ion intensity maps displaying the spatial distribution for statistically significant mass features were created using R. Before generating the ion maps, the intensity values for the selected mass features were normalized to the maximum intensity within the image, measured for each mass value individually. Venn diagrams displaying unique and common masses amongst different treatments were drawn using the jvenn tool (64).

**Direct Analysis in Real Time Mass Spectrometry (DART-MS) data analysis**

DART-MS data analysis was performed with Xcalibur 2.2 (Thermo) and files were exported as mzdata files and further analyzed using MZmine V2.24 (Copyright © 2005-2015 MZmine Development Team) (12). Compounds were identified via their mass spectra using the online KEGG database (14, 15). After deconvolution and mass identification, peak lists containing the mass features of each treatment were exported in csv file format and uploaded to a local copy of Metaboanalyst V3.0 (www.metaboanalyst.ca) running on a local UNIX server for statistical analysis (13). To identify significant abundant masses ONE-WAY-ANOVA with post-hoc TUKEY test was performed between the data sets. Masses were considered to be statistical relevant if p- and FDR- values were $\leq 0.05$.

#### References Supplementary Material


1. Garbeva P, van Elsas JD, de Boer W. 2012. Draft Genome Sequence of the Antagonistic Rhizosphere Bacterium Serratia plymuthica Strain PRI-2C. Journal of Bacteriology 194:4119-4120.
2. Tyc O, de Jager VCL, van den Berg M, Gerards S, Janssens TKS, Zaagman N, Kai M, Svatos A, Zweers H, Hordijk C, Besselink H, de Boer W, Garbeva P. 2017. Exploring bacterial interspecific interactions for discovery of novel antimicrobial compounds. Microbial Biotechnology 10:910-925.
3. de Boer W, Wagenaar AM, Klein Gunnewiek PJA, van Veen JA. 2007. In vitro suppression of fungi caused by combinations of apparently non-antagonistic soil bacteria. Fems Microbiology Ecology 59:177-185.
4. Tyc O, van den Berg M, Gerards S, van Veen JA, Raaijmakers JM, de Boer W, Garbeva P. 2014. Impact of interspecific interactions on antimicrobial activity among soil bacteria. Frontiers in Microbiology 5,.
5. Garbeva P, de Boer W. 2009. Inter-specific Interactions Between Carbon-limited Soil Bacteria Affect Behavior and Gene Expression. Microbial Ecology 58:36-46.
6. Lane DJ. 1991. 16S/23S rRNA sequencing. In: Nucleic acid techniques in bacterial systematics. John Wiley and Sons, New York, NY.
7. Edwards U, Rogall T, Blocker H, Emde M, Bottger EC. 1989. Isolation and Direct Complete Nucleotide Determination of Entire Genes - Characterization of a Gene Coding for 16s-Ribosomal Rna. Nucleic Acids Research 17:7843-7853.
8. Kanehisa M, Sato Y, Morishima K. 2016. BlastKOALA and GhostKOALA: KEGG Tools for Functional Characterization of Genome and Metagenome Sequences. J Mol Biol 428:726-731.
9. Sayers EW, Cavanaugh M, Clark K, Ostell J, Pruitt KD, Karsch-Mizrachi I. 2018. GenBank. Nucleic Acids Research 47:D94-D99.
10. Garbeva P, Hordijk C, Gerards S, de Boer W. 2014. Volatiles produced by the mycophagous soil bacterium Collimonas. Fems Microbiology Ecology 87.
11. Tyc O, Zweers H, De Boer W, Garbeva P. 2015. Volatiles in inter-specific bacterial interactions. Frontiers in Microbiology 6.
12. Pluskal T, Castillo S, Villar-Briones A, Oresic M. 2010. MZmine 2: modular framework for processing, visualizing, and analyzing mass spectrometry-based molecular profile data. Bmc Bioinformatics 11:395.
13. Xia J, Sinelnikov IV, Han B, Wishart DS. 2015. MetaboAnalyst 3.0-making metabolomics more meaningful. Nucleic Acids Research 43:W251-7.
14. Anonymous. The KEGG Database, 'In Silico' Simulation of Biological Processes doi:10.1002/0470857897.ch8.
15. Kanehisa M. 2008. The KEGG Database. *In* Novartis Foundation GBaJAG (ed), The KEGG Database In 'In Silico' Simulation of Biological Processes doi:10.1002/0470857897.ch8.
